# Multiscale simulations examining glycan shield effects on drug binding to influenza neuraminidase

**DOI:** 10.1101/2020.08.12.248690

**Authors:** Christian Seitz, Lorenzo Casalino, Robert Konecny, Gary Huber, Rommie E. Amaro, J. Andrew McCammon

**Author notes:** Co-corresponding author: Christian Seitz, Twitter handle: @chem_christian, Co-corresponding author: Rommie E. Amaro, Twitter handle: @RommieAmaro.

## Abstract

Influenza neuraminidase is an important drug target. Glycans are present on neuraminidase, and are generally considered to inhibit antibody binding via their glycan shield. In this work we studied the effect of glycans on the binding kinetics of antiviral drugs to the influenza neuraminidase. We created all-atom *in silico* systems of influenza neuraminidase with experimentally-derived glycoprofiles consisting of four systems with different glycan conformations and one system without glycans. Using Brownian dynamics simulations, we observe a two- to eight-fold decrease in the rate of ligand binding to the primary binding site of neuraminidase due to the presence of glycans. These glycans are capable of covering much of the surface area of neuraminidase, and the ligand binding inhibition is derived from glycans sterically occluding the primary binding site on a neighboring monomer. Our work also indicates that drugs preferentially bind to the primary binding site (i.e. the active site) over the secondary binding site, and we propose a binding mechanism illustrating this. These results help illuminate the complex interplay between glycans and ligand binding on the influenza membrane protein neuraminidase.

**Statement of Significance:** The influenza glycoprotein neuraminidase is the target for three FDA-approved influenza drugs in the US. However, drug resistance and low drug effectiveness merits further drug development towards neuraminidase, which is hindered by our limited understanding of glycan effects on ligand binding. Generally, drug developers do not include glycans in their development pipelines. Here, we show that even though glycans can reduce drug binding towards neuraminidase, we recommend future drug development work to focus on strong binders with a long lifetime. Furthermore, we examine the binding competition between the primary and secondary binding sites on neuraminidase, leading us to propose a new, to the best of our knowledge, multivalent binding mechanism.

## Introduction

It has been long appreciated that glycans on influenza membrane proteins help shield the virus from the host immune system’s antibodies (1–7). Unrecognized glycosylation differences can also attenuate influenza vaccines (8). In one study, glycans were shown to reduce epitope accessibility and drug binding to receptor proteins (9). Glycans can clearly influence antibody binding due to their presence in the antibody binding site. However, it remains to be seen whether this glycan shielding and glycoprofile variability is also a concern for influenza drugs, recognizing that these drugs are smaller than human antibodies, and the fact that glycans present themselves near, but not directly inside, the catalytic sites. Currently there are three FDA-approved influenza neuraminidase (NA) antivirals in the US: Tamiflu (oseltamivir), Relenza (zanamivir) and Rapivab (peramivir), all of which have lingering questions over their efficacy, side effects, and drug resistance (10, 11). This necessitates the need for further drug development against influenza (12).

Drug developers have many hurdles to clear when designing a new influenza drug: classical ADMET characteristics, clinical trials and governmental regulations, among others. What is not often considered is the viral glycosylation state. The glycosylation state is the assemblage of glycans, linkages of sugars found on the surface of about half of all proteins (13). Influenza contains N-linked glycosylation sites, defined by the Asn-X-Ser/Thr sequon, where X can be anything besides proline (14). This leads to the so-called glycan shield, where glycans on the protein surface are capable of accessing much of the protein’s surface area, and potentially shielding it from outside interactions (15–22). Among their many biological functions, glycans play a crucial, but complex role in viral infection (23). One salient example of glycan function in influenza is how they help the virus evade the immune system (1–7). Furthermore, glycans are capable of affecting receptor binding in influenza (5, 7, 24–27).

Traditionally, glycans have been difficult to study due to their flexibility and heterogeneity. Most of the glycan characterization studies are done through mass spectrometry, which can yield highly variable glycoprofile data, such as differences in the degree of post-translational modifications, sequon occupancy, and type of glycan, for different strains of influenza (28–33). Similarly, glycan occupancy levels are not consistent across studies, even when using the same cell line and strain of influenza (34, 35) These discrepancies may arise from differences in system setup, sample preparation, cell culturing and/or analysis method, which increases the difficulty in determining the transferability of experimental glycan results. Though not well understood, the number and position of the glycosylation sites on influenza can change over time as a result of antigenic drift (36–39). This increases the glycoprofile variability, effectuating irregular but significant changes in the glycan shield over time.

Considering the variability and immune evasion function of the glycan shield, it remains to be seen what effect this shield has on small-molecule antiviral drug binding to viral surface proteins. Previous work has shown that, depending on the viral strain and receptor mimetic used, removing viral glycans can improve binding to cell receptor mimetics (40–42). Other studies have shown that these viral glycans decrease binding of other cell receptor mimetics (27, 43–46). Regardless, antiviral drugs will be much smaller than a receptor mimetic, and it is not clear whether this size difference means antiviral drugs will still be affected by the viral glycans. An earlier study by Kasson and Pande, using 100 ns molecular dynamics (MD) simulations, showed reduced binding of α2-3-sialyllactose trisaccharides to hemagglutinin due to glycans (43). A recent review concluded that the viral glycosylation state should be considered when designing small molecule antivirals (47).

Focusing on how small molecule antivirals are affected by the glycan shield, we combine results from distinct BD and MD simulations into an integrated multiscale simulation study. We have utilized BD to estimate the rates of binding of small molecules to the primary (i.e. active/catalytic) and secondary (i.e. hemadsorption) binding sites of influenza neuraminidase in glycosylated and unglycosylated states. We see that the glycan shield is capable of moderately inhibiting drug association to the primary binding site of NA on the order of two to eight times. Small molecule association is faster to the primary binding site than the secondary binding site. Ligand binding is independent between the primary and secondary sites – the presence of one site does not influence binding at the other site. Overall, this work provides insights into the impact of glycans on small-molecule binding to NA.

## Methods

In this study, we use Brownian dynamics (BD), which has been previously used to simulate protein-small molecule association (48–51). Specifically, it has also been used to simulate the association of small molecules to influenza neuraminidase (52–54). BD makes the implicit assumption that long-range electrostatics and stochastic collisions with solvent molecules are the driving forces behind protein-ligand binding (55). Therefore, it is an efficient method to simplify binding to describe electrostatically-influenced diffusion. Using BD allows for a reduction in system complexity and a focus on specific modulations of ligand association.

To assess whether glycans affect small molecule binding to NA, we created an *in silico* NA model using the strain of influenza A virus, A/Viet Nam/ 1203/2004(H5N1) and tetrameric PDB 2HTY, with Uniprot ID Q6DPL2 (56). Building on this structure, we generated five NA constructs: (i) one unglycosylated model; (ii) one glycosylated model with web server-derived glycan conformations; (iii) three glycosylated models, each with unique, biologically-relevant glycan conformations derived from all-atom MD simulations that were based on (ii) as the starting structure. Finally, we ran BD simulations using these models to examine binding characteristics of oseltamivir, zanamivir and sialic acid. To note, the BD input and results files are provided on Github (https://github.com/cgseitz).

### Setup of the unglycosylated model

The unglycosylated model was built using an avian H5N1 strain and was used as a basis for the other models. We picked this strain of influenza because it contains a glycosylation site at N146, a member of the 150-loop that hangs over the primary binding site, as shown in **Figure 1**. This close proximity provides a good test of whether glycans were capable of interfering with ligand association to influenza neuraminidase. As the BD simulations used here keep bonds rigid, it was necessary to select ligand conformations that represented a bound state and protein conformations that represented an open state, to properly approximate the initial binding contact. Thus, we selected a crystallized apo head region of the strain mentioned above (PDB 2HTY) (56). The stalk region has not been crystallized for any influenza NA and is unlikely to influence ligand association due to its large distance from the distal binding sites, so it was not modelled. The crystallized calcium ions were retained throughout, while the crystallized glycan fragments were removed (57).

**Figure 1.**
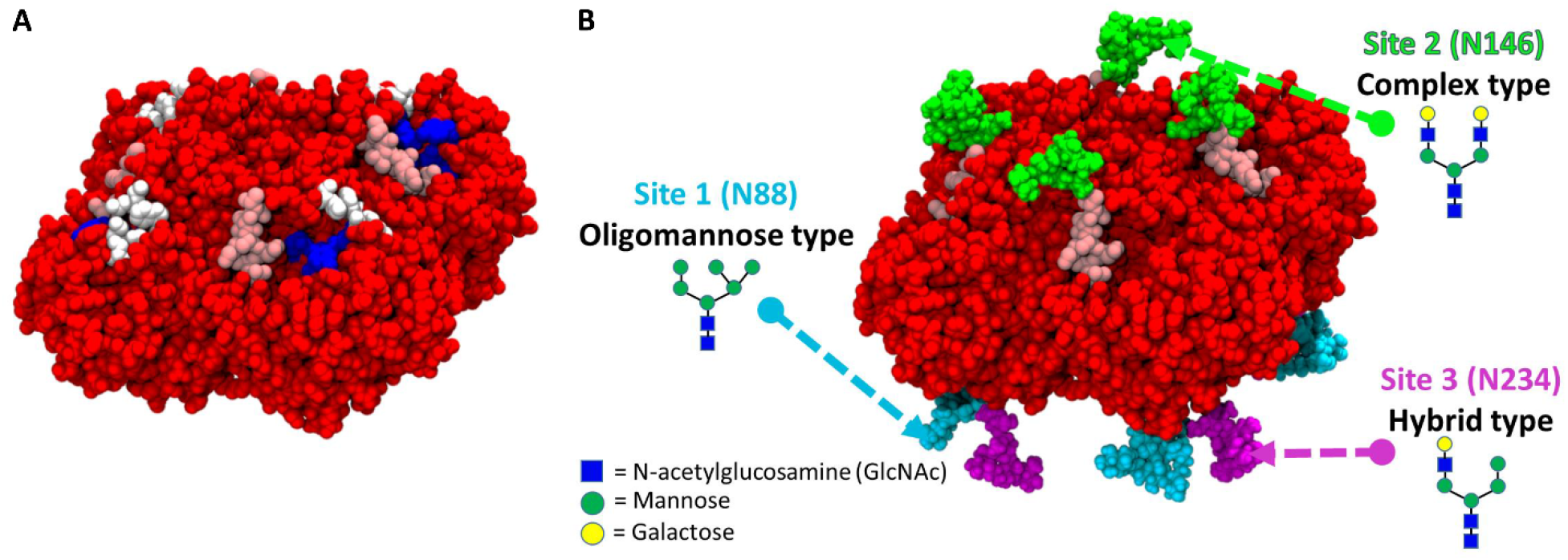
Structure of the NA head. (A) The relation of the binding sites to the 150 loop and each other are shown. The NA protein is in red, the 150-loop is in pink, the primary binding site is in blue, and the secondary binding site in white. (B) The glycan structures and types used in this study are shown in relation to the 150 loop. The glycan on top of each monomer is attached to the 150 loop. Three unique glycan structures and types were used in the simulations. The font color of the glycosylation site refers to the corresponding glycan structure on the protein.

2HTY was crystallized with a Y171H mutation (PDB numbering), which was reversed for this project through PyMOL (58). The histidine rotamer was chosen to be the one with the highest occurrence in proteins, according to PyMOL. The crystal structure contained a broken backbone between P169 and N170 which was fixed through Schrodinger Maestro; subsequently residues 168-171 (on each side of the fixed bond) were minimized through Maestro (59). The same procedure was done for the broken backbone between V411 and Q412: the bond was created and residues 410-413 were minimized. This refitting was done for each monomer in the tetramer. The pH was set to pH 6.4, as this was done in the reference k_on_ experiments (60, 61). Using this pH, protonation states on the neuraminidase were assigned using PROPKA (62). The protonation assignments were done through the PDB2PQR server (63). Partial charges on the protein were assigned according to the AMBER99 force field (64). Parameterizing the glycans needed special treatment as there are not glycan parameters in the AMBER99 force field. We used the GLYCAM_06h-1 parameters as these would be consistent with the AMBER99 force field (65).

### Setup of the glycosylated NA model with web server-derived glycan conformations

To build the first glycosylated construct (with web server-derived glycan conformations), the unglycosylated NA structure was uploaded to the Glyprot server, and three representative glycans were added to each NA monomer, for a total of 12 glycans on the NA homotetramer (66). Though there is experimental variability in glycosylation site occupancy, we decided to place a glycan in each glycosylation site to see the maximum potential effect the glycoprofile can have on ligand association. Considering most of the human H5N1 transmission came directly from avian sources, the glycans used to model this structure came from an avian (hen egg) source for growing these glycans (30). Additionally, this dataset is the only one containing structures experimentally found on influenza NA (30). We chose representative glycans from this dataset, however we note that both larger and smaller glycans will exist in nature; these size differences may slightly affect the results presented here. The exact glycans were selected as shown in **Table 1**.

**Table 1.** Glycan structures from the Glyprot web server. The “Glycan structure” entries came from experimental results (30). These structures consist of N-acetylglucosamine (GlcNAc), mannose (Man), N-acetylhexosamine (HexNAc) and hexose (Hex). HexNAc and Hex were interpreted according to their corresponding “Glyprot identifier” and the structures shown in **Figure 1**.

| # | Glycosylation site | Glycan structure | Glyprot identifier |
| --- | --- | --- | --- |
| 1 | N88 | (GlcNAc)<sub>2</sub>(Man)<sub>3</sub>(Hex)<sub>3</sub> | gly_8792.pdb |
| 2 | N146 | (GlcNAc)<sub>2</sub>(Man)<sub>3</sub>(HexNAc)<sub>2</sub>(Hex)<sub>2</sub> | gly_9196.pdb |
| 3 | N234 | (GlcNAc)<sub>2</sub>(Man)<sub>3</sub>(HexNAc)<sub>1</sub>(Hex)<sub>2</sub> | gly_8582.pdb |

To better diversify our system, three glycosylation sites (termed as site #1, site #2 and site #3) present on each monomer were linked to three different glycan types. Importantly, the four monomers (termed as monomer A, monomer B, monomer C, and monomer D) of our homotetrameric NA model were symmetrically glycosylated, meaning that sites #1, #2 and #3 were populated with the same glycan across monomers.

This resulted in glycans A1-3, B1-3, C1-3, and D1-3, where all glycans linked at site #1 have identical structures (but not necessarily identical conformations), all glycans linked at site #2 have identical structures, and all glycans linked at site #3 glycans have identical structures. The glycans selected are shown in **Figure 1**.

### Molecular dynamics simulations

Starting with the structure containing web server-derived glycan conformations, MD was then used to generate representative glycan conformations, with the assumption that MD would provide realistic conformations of glycans within a microsecond’s worth of sampling (67, 68). The first step was porting the structure with web server-derived glycans into CHARMM-GUI to prepare the structure for MD (69–74). The disulfide bonds were taken from Uniprot ID Q6DPL2. The system was embedded into a box described with explicit water molecules using the TIP3P model (75). An ion model was used as described previously (76). The full system had a size of 299,732 atoms. An ionic solution of 0.15 M NaCl was used, and the CHARMM36 all-atom additive force fields were used for the protein and the glycans (77). Molecular dynamics simulations were run using GPU-accelerated AMBER18 with an NPT ensemble (78, 79). The system was initially minimized for a total of 5000 cycles using a combination of steepest descent and conjugant gradient methods (78, 79). Equilibration in an NPT ensemble was performed for 125 ps, using a timestep of 2 fs and the SHAKE algorithm to constrain all bonds involving hydrogen (80). The equilibration temperature was set at 298 K and regulated through a Langevin dynamics thermostat (81, 82). The pressure was fixed at 1 bar through a Monte Carlo barostat (83). These simulations were run using Extreme Science and Engineering Discovery Environment (XSEDE), specifically the Comet supercomputer housed at the San Diego Supercomputer Center (84). Periodic boundary conditions were used with a non-bonded short-range interaction cutoff of 12 Å and force-based switching at 10 Å. Particle mesh Ewald was used for the long-range electrostatic interactions (85). For the production runs, the temperature was set at 310.15 K (61). After equilibration, this system was cloned into 50 identical replicates. Each one was run in parallel for 20 ns each with a unique starting velocity, totaling 1 μs of sampling.

### Glycan clustering and setup of the glycosylated NA models with representative glycan conformations

Once the MD simulations finished, the trajectory of each glycan was concatenated independently of the rest of the system. Each of these individual glycan trajectories were then clustered using GROMACS-based GROMOS clustering with an RMSD cutoff of 2.5 Å (86). This number was chosen so the three most populated clusters would represent at least 50% of the total glycan conformations in each of the simulations. The central structure, defined as the structure with the smallest average RMSD from all other members of the cluster, from the top cluster of each of the 12 glycans was then selected; the pyranose ring from the reducing end of the glycan was then aligned to the analogous pyranose ring of the corresponding glycan from the Glyprot-glycosylated structure as this should be the most stable part of the glycan (87). The Glyprot glycans were removed and the glycans from the MD simulations were attached through Schrodinger Maestro, to create a new NA system with each glycosylation site inhabited by the central structure of the most representative conformation from the MD simulations. This was then repeated for the second and third most representative glycan clusters from the MD simulations.

### Ligand setup

The sialic acid structures used were drawn from PDB 1MWE, which crystallized the boat conformation in the active site and the chair conformation in the secondary site (88). The chair conformation of sialic acid was crystallized with a missing carboxylate group, which was added through Schrodinger Maestro to model an energetically-favorable gauche conformation. Zanamivir was extracted from the 3CKZ crystal structure (60). Oseltamivir was extracted from the 3CL0 crystal structure (60). A 2D comparison of these ligands can be seen in **Figure S1**, showing their structural similarities; we note that all mentions in this study of oseltamivir pertain to Tamiflu’s active metabolite oseltamivir carboxylate. These ligands were then uploaded to the PRODRG server to add hydrogens (89). Charges according to the AMBER99 force field were added through the PDB2PQR server (63, 64).

### Brownian dynamics (binding pairs and simulations)

BD simulations were run using Browndye (90). Even though the MD was run with the AMBER18 force field and the BD was run with the AMBER99 force field, we assume these to be sufficiently independent steps and the slight force field differences should not appreciably affect the results, especially as our k_on_ numbers are relative, not absolute. The charges for the protein and ligands were reassigned according to the AMBER99 force field (64). The temperature was set to 310.15 K, which was the temperature for the referenced k_on_ experiments (60, 61). The ions used are shown in **Table S1**. These ions were selected to mimic the ion and buffer concentration of the reference k_on_ experiments (60, 61).

The experimental assay used 5 mM CaCl_2_ and 32.5 mM MES buffer (60). The Ca^2+^ and Cl^−^ concentrations were simply calculated by finding their ionic strengths. MES buffer is prepared with Na^+^; the concentrations of the buffer and Na^+^ at pH 6.4 were calculated with the Henderson-Hasselbalch equation (91, 92). This resulted in an overall ionic strength of 0.039 M. The calcium, chlorine and sodium van der Waals radii were taken from the literature (64, 93). The MES radius was determined by building it in Schrodinger Maestro and measuring it in VMD (94).

APBS was used to create the electrostatic grids needed by Browndye for these simulations (95). The grid spacings are listed in **Table S2**.

The solvent dielectric was set to 78 while the protein dielectric was set to 4. Desolvation forces were turned off. The Debye length, determined from the concentration and charges of the ions in the solution, was set to 15.7 Å. In Browndye, the b radius is defined as the starting radius for the ligand trajectories, at a distance where the force between the protein and ligand is independent of orientation. This distance is determined from the hydrodynamic center of the receptor. Because of the different glycan conformations used, the b radius differed slightly between systems. If a ligand reaches what is known as the q radius, the trajectory either ends as a non-association or is restarted from the b radius according to Browndye’s algorithm. The q radius is defined as 1.1 times the b radius distance. The b radius ranged between 109 Å and 112 Å depending on the system, and the q radius ranged from 120 Å to 123 Å. The exact b and q radius values for each system are shown in **Table S3**.

BD simulations were run on all five NA models generated (i.e. unglycosylated, glycosylated with web server-derived glycans, and the three systems with MD-derived glycan conformations). These simulations totaled 10 million trajectories for each ligand/binding site pair, consistently giving reproducible rates within the small level of error reported and resulting in 600 million trajectories total. Reproducible rates will be obtained by having a binding probability of around one in a million trajectories; we found we could roughly obtain these probabilities by using 10 million trajectories for each ligand/binding site as has been reported previously (53). This number of trajectories produced error values comparable to those seen in the reference experimental studies, as seen in **Figure S2**. For systems where we saw at least one binding event, the number of binding events ranged from two to 889 (see Supporting Material for details).

### Brownian dynamics reaction criteria

BD simulations using Browndye requires the creation of reaction criteria, consisting of a list of protein-ligand atom pairs and a cutoff distance. If any three of these pairs simultaneously came closer than the cutoff distance, we assume the ligand will associate. The cutoff distance was empirically determined to be 3.228 Å; this distance approximately yielded the experimental k_on_ rates for both oseltamivir and zanamivir (60). There are no other experimental k_on_ rates towards the primary site of H5N1, and no referenced rates at all for the secondary site. The referenced k_on_ experiments were done with glycans attached to NA and measured to the full tetramer; this was confirmed in personal correspondence with the corresponding author (Stephen Martin of the MRC National Institute for Medical Research, correspondence on July 21, 2018). Considering that the reaction criteria and reaction distance were created for oseltamivir and no significant changes were made before applying them to zanamivir, we can safely assume that they are generalizable for sialic acid, an analog of both oseltamivir and zanamivir (**Figure S1**).

The protein-ligand atom pairs were taken from crystal structures of ligands in the primary and secondary sites of neuraminidase for each monomer, and simulations were run for the full tetramer. The primary binding site was determined according to the crystallized binding pocket for our strain of neuraminidase (56). This pocket is noted to have a surface area of 941.3 Å^2^ and a volume of 574.8 Å^3^ (96). The secondary site contacts were determined from a structure of influenza A/tern/Australia/G70C/75 (88). However, all the secondary site residues are conserved between that strain and the strain used in our simulations. The combined site simulations are defined as simulations with criteria allowing for association to either the primary or secondary site; it is simply a simulation run with a concatenation of the binding criteria for these sites.

In this work, we define binding site contacts to be those protein-ligand contacts seen in crystal structures. From these contacts, we created protein-ligand atom pairs in Browndye to determine when a reaction has occurred in our BD trajectories. There are seven primary binding site contacts reported between oseltamivir and the 3CL0 crystal structure (60). These binding site contacts are reported in **Table S4** and **Figure S3**. There are five primary binding site contacts reported between sialic acid and the 1MWE crystal structure; all five of these are analogous to those seen for oseltamivir (88). The binding site contacts for sialic acid are registered in **Table S5** and **Figure S4**. There is one primary binding site contact reported between zanamivir and the 3CKZ crystal structure; this one is analogous to one seen in oseltamivir (60). The binding site contacts from oseltamivir were transferred to zanamivir retaining the one contact seen in the 3CKZ crystal structure and are reported in **Table S6** and **Figure S5**. Using the structural similarities of sialic acid and zanamivir to oseltamivir, analogous primary binding site atom pairs were created so that each ligand had seven primary protein-ligand atom pairs.

There are five secondary binding site contacts reported between sialic acid and the 1MWE crystal structure (88). These contacts are reported in **Table S7**. There are no published reports of crystal structures of oseltamivir or zanamivir in the secondary binding site, so five analogous secondary binding site protein-ligand atom pairs were created for oseltamivir (**Table S8**) and zanamivir (**Table S9**) to match those seen in sialic acid, so that each ligand had five secondary binding site protein-ligand atom pairs.

## Results

### Glycan shield and individual glycan clusters

To pare down the data from 1 μs of cumulative MD sampling and pick out biologically-relevant glycan conformations, we clustered each glycan from the MD simulations. The glycan trajectories were extracted and affixed on the static NA crystal structure, to reveal the conformational space they can access (**Figure 2**). Visualizing these glycan trajectories on the NA structure gives a qualitative representation of how much volume and surface area the glycans are capable of accessing. Keeping in mind the primary and secondary binding sites are located just beneath the glycans (**Figure 1**), the size and flexibility of the glycans here shows that they have the capability to “shield” the binding sites from ligand association.

**Figure 2.**
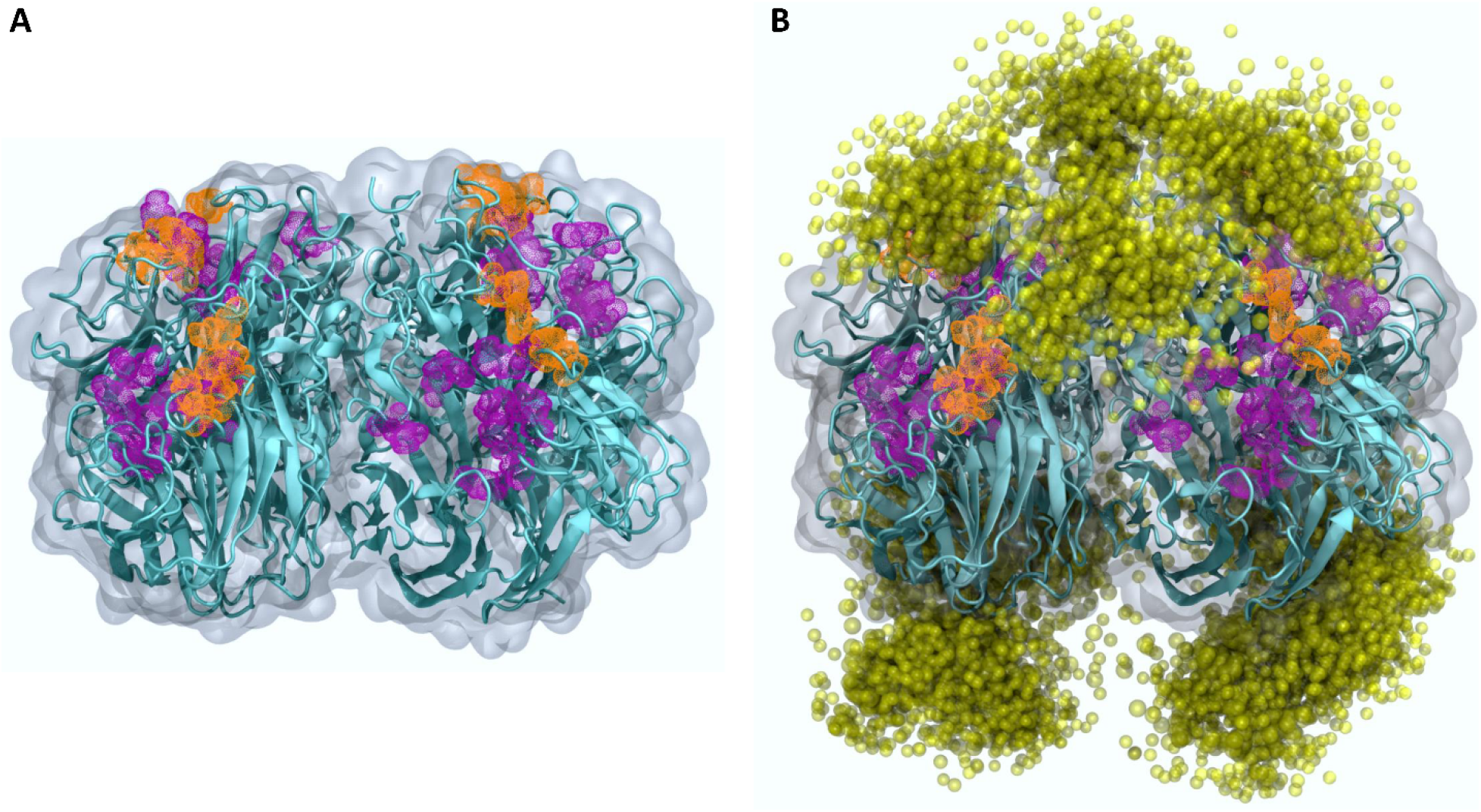
Glycan shielding on NA. (A) Molecular representation of the unglycosylated NA’s homotetrameric head showing accessible binding sites. The static structure of NA is in teal. The primary binding sites are in purple, and the secondary binding sites are in orange. (B) Molecular representation of the glycosylated NA’s homotetrameric head showing much of its surface area shielded. The static structure of NA is in teal, the primary binding sites are in purple, and the secondary binding sites are in orange. The glycan shielding density is represented with a yellow cloud of spheres using multiple layered glycan conformations from the MD simulations. Each sphere approximates a monosaccharide. A total of 10 frames, using a stride of 100, were selected from the 1000 frames-long trajectory obtained upon concatenation of all the MD simulations.

The three most representative clusters for each glycan were extracted from the MD simulations. The central structure from each cluster was compared with the conformation generated from Glyprot. These clusters show some conformational diversity, but none show a particularly similar conformation to the Glyprot structure. However, the third glycan in each monomer shows a markedly decreased conformational diversity compared to the other two monomers. The clustering results from each monomer show the same trends; the results from monomer A are shown in **Figure 3**, while the results from monomer B (**Figure S6**), monomer C (**Figure S7**), and monomer D (**Figure S8**) are shown in the Supporting Material.

**Figure 3.**
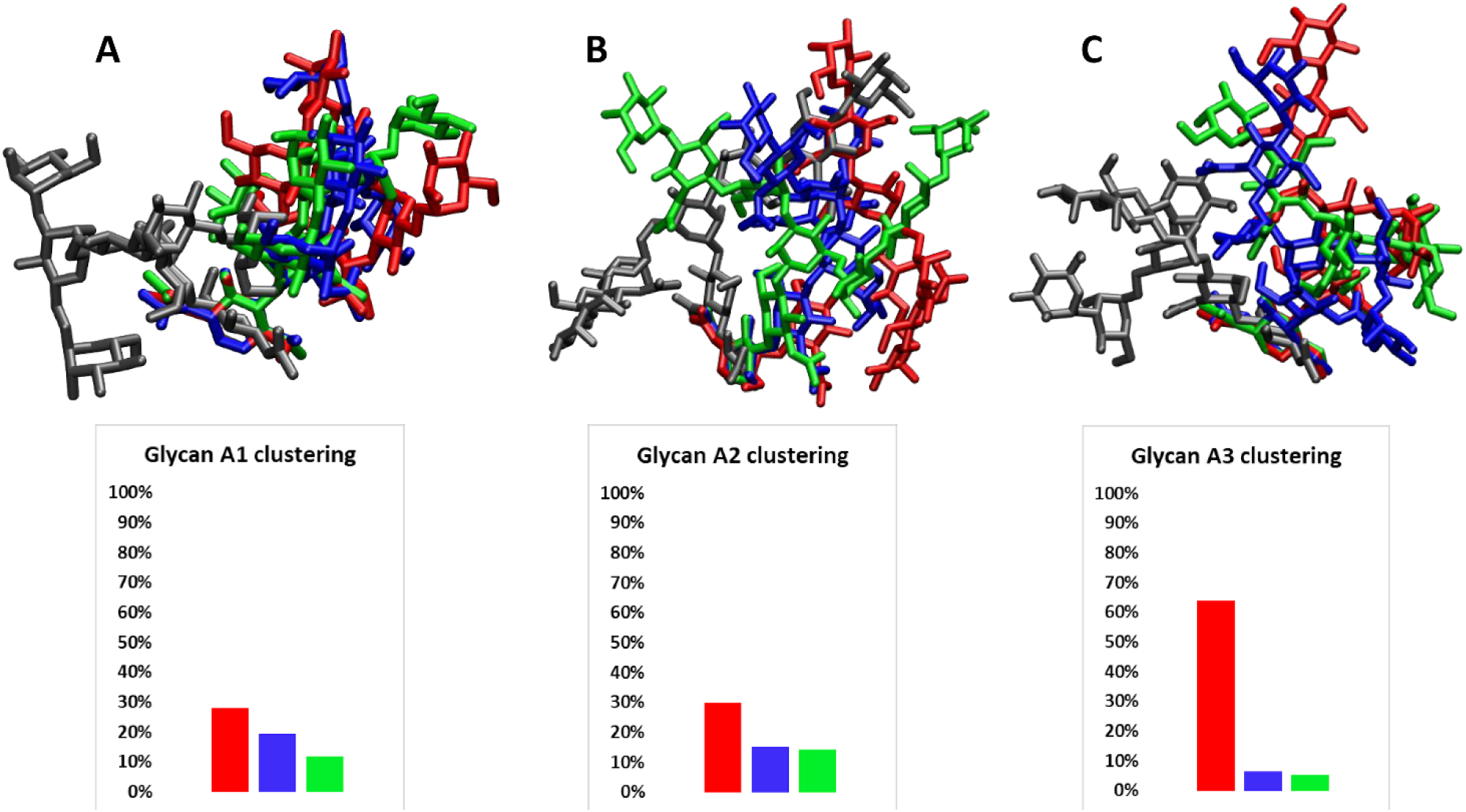
Clustered glycans on monomer A. The glyprot structure is in gray, while the other colors represent clusters from the MD simulations.

The glycans on the top of the NA head are all situated directly over the primary binding site on their own monomer. Conformations extracted from the MD simulations universally bent towards the secondary binding site located on the adjacent monomer, leaving the primary binding site closest to them accessible (**Figure 4**).

**Figure 4.**
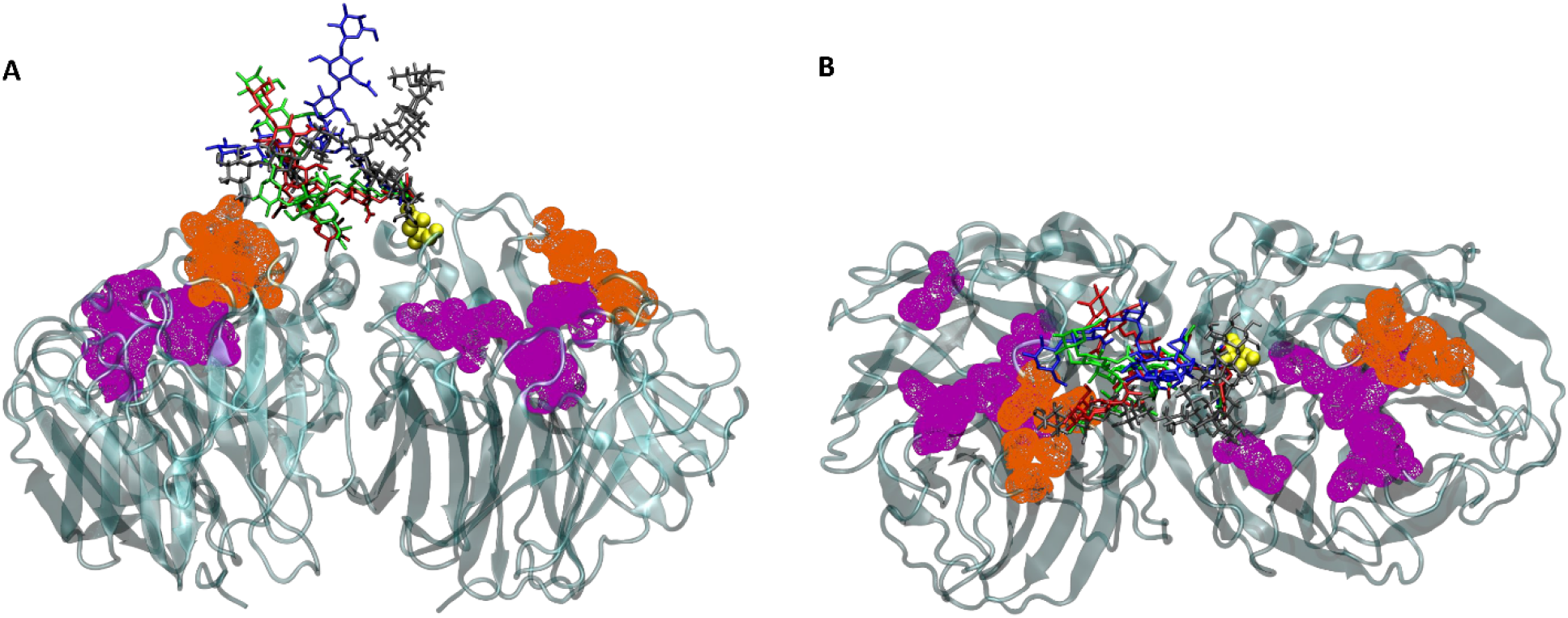
The superposition of each glycan onto the static NA structure shows the conformational variation of the glycans. (A) Side view and (B) top view of the glycosylated NA model with MD-derived glycan conformations. For ease of viewing, only two monomers are shown here, with the glycans only coming from the monomer on the right in panel (A) and panel (B). The yellow glycosylation site N88 and the attached glycans reside in the same monomer. The glycans bend away from the binding sites on their monomer towards the binding sites on the neighboring monomer. This is seen for each monomer. The primary binding sites are in purple and the secondary binding sites are in orange. The linkage between the glycans and the protein is in yellow. The NA structure is in teal. The Glyprot conformation is in gray, the first conformation from the MD simulations is in orange, the second conformation is in blue, and the third conformation is in green.

### Association rates of oseltamivir, zanamivir and sialic acid

To be confident in our computed association rates, we first needed to benchmark our system against experimental results. We created empirically-derived system criteria for the association of oseltamivir to the primary binding site of glycosylated NA, as described in the methods. After matching the experimental association rate with oseltamivir, the same parameters were applied to zanamivir. These are the only two experimental association rates for H5N1 NA.

Subsequently, we investigated the association of oseltamivir and zanamivir to the primary sites of glycosylated NA, obtaining association rates of 2.52 ± 0.21 /μM·s for oseltamivir and 0.47 ± 0.09 /μM·s for zanamivir. These are in agreement with the experimentally-measured rates of 2.52 ± 0.21 /μM·s and 0.95 ± 0.21 /μM·s, respectively, as visualized in **Figure S2** (60). Considering the experimental systems were glycosylated, we had to pick one glycan conformation to use for computing these benchmarks in our glycosylated system; for reproducibility we chose the conformation generated from the Glyprot server. We note that choosing a different conformation for our computed benchmark would change the absolute association rates by a scaling factor, but the trends would remain the same.

Since the predicted k_on_ for oseltamivir and zanamivir both matched up well with the experimental rates, the system proved to be transferable to ligand analogs for the primary site. We then applied the same criteria to two different conformations of sialic acid, boat and chair, to probe if the association rate was dependent on conformation. This was done in addition to analyzing how association rate was modulated by different functional groups, via comparisons of ligand analogs such as oseltamivir, zanamivir and sialic acid.

With the binding criteria set up, we calculated the association rates of each of the ligands to the primary site (**Figure 5A**). These results show two important findings. First, there is not a large difference in association rates between the system with Glyprot glycans and the unglycosylated system. This shows that a glycan may adopt a conformation where it does not inhibit ligand binding much at all. The second finding is that the glycans from the MD simulations all show a moderate level of inhibition, more than the system with Glyprot glycans. This shows that biologically-relevant glycan conformations will likely exhibit a moderate level of inhibition towards ligand binding. Combining the first and second finding discussed in this paragraph, glycans are capable of perturbing ligand binding to NA.

**Figure 5.**
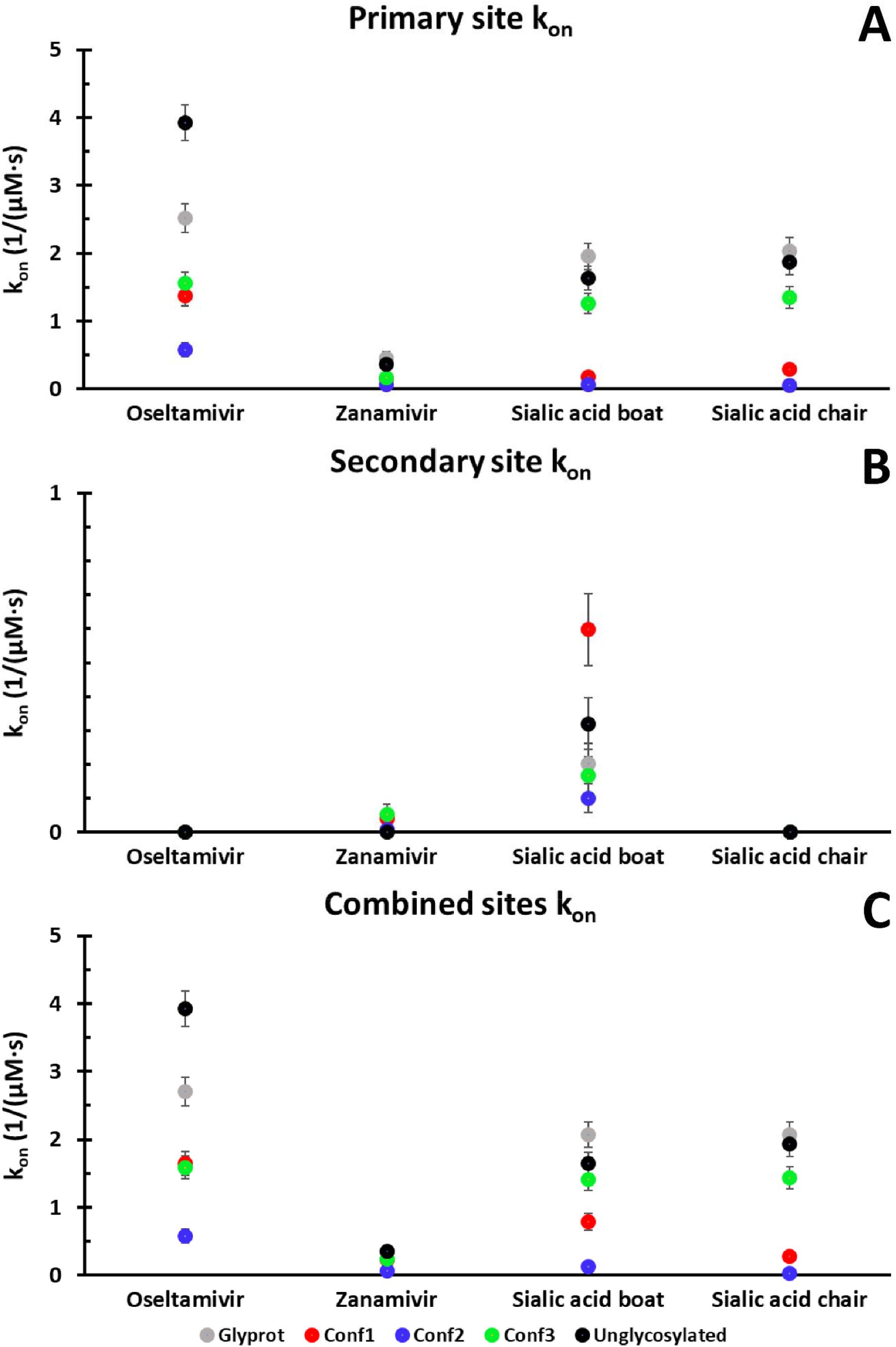
Association rates of each ligand to the primary and secondary sites. Conf1 is the glycan structure from the most populated cluster from the MD simulations. Conf2 is from the second most populated cluster, and conf3 is from the third most populated cluster. The association rates using glycans structures downloaded from Glyprot are shown in gray. The association rates using structures derived from the MD simulations are in bright, colorful shades whereas the others are in grayscale. The association rates without using any glycans are shown in black. (A) The glycan structures from the MD simulations show a moderate association rate inhibition to the primary binding site irrespective of ligand chosen. (B) Little association is seen to the secondary binding site. Note the different y-axis used to be able to see the small amount of binding. (C) Association rates of trajectories run with either the primary site or secondary site as the trajectory end point. Similar to (A), the glycans structures from the MD simulations in (C) show a moderate inhibition of ligand association. The raw data for this figure is seen in Table S10 (oseltamivir), Table S11 (zanamivir), Table S12 (sialic acid boat conformation), and Table S13 (sialic acid chair conformation).

There are no experimental association rates for ligands to the secondary site, so criteria were chosen based off of crystal structure data and discussed in the methods. Only sialic acid has been crystallized in the secondary site of avian NA, so binding site criteria for the secondary site were extracted from that structure and used to create the criteria for oseltamivir and zanamivir, as discussed in the methods (88). A previous BD study suggested that oseltamivir can bind to the avian NA secondary site (52). A follow-up NMR study also suggested that the oseltamivir binds to the avian NA secondary site (97). However, a more recent experimental study disagreed with these findings and did not see oseltamivir binding to the avian NA secondary site (98). Considering the disagreement with oseltamivir binding to the secondary site, we decided to test this and secondary site binding for zanamivir as well. The computed association rates towards the secondary site show a markedly different story than those to the primary site (**Figure 5B**). None of the ligands exhibited noticeable binding towards the secondary site, with the exception of the boat conformation of sialic acid. Even with this conformation, there is no consistent trend when compared to primary site binding. Although the boat conformation sialic acid displays a small amount of binding, the chair conformation does not show binding. These results show that we can differentiate between these two sialic acid conformations at the BD level of theory.

Finally, trajectories were run where the ligand could associate to either the primary site or the secondary site (**Figure 5C**). Intriguingly, the results are essentiallly a concatenation of the rates seen for the primary and secondary sites individually. Considering the low level of secondary site binding, the trends here are the same as seen for the primary site.

As can be seen in **Figure S1**, there is a formal charge difference between the ligands: Sialic acid contains a formal charge of −1 while oseltamivir and zanamivir are neutral. Running test BD trajectories without charge treatment (results not shown), we saw analogous results to those seen in **Figure 5**. This meant that only the sterics of the systems affected binding, not electrostatics. Clearly one or a few of the structural differences between the ligands play outsized roles in affecting the association rates. In this work we did not further probe which exact atoms in the ligands will change the association rates.

## Discussion

### The glycan conformational flexibility underpinning the glycan shield

Biologically, the influenza replication cycle is propagated through NA recognizing and cleaving sialic acid. This study compares the interplay between that molecular recognition process and NA’s aforementioned glycan shielding capabilities. This interplay is simplified here by approximating ligand binding as a diffusion-governed association process, modulated by protein electrostatics.

Previous studies have shown that viral proteins can exhibit a degree of glycosylation large enough to partially protect a variety of viruses from immune system antibodies; this is termed the viral glycan shield (18, 21, 99–101). From static structures one can envision the shielding that glycans can provide, but a dynamic representation better depicts the steric barrier encountered by immune system antibodies and drugs (102). In our single NA protein, we see that glycans are capable of covering most of the NA surface area, as shown in **Figure 2**. This is consistent with studies explaining how the influenza glycan shield can cloak the influenza virion from the immune system (5–7). The glycans can access a large volume, allowing for a considerable shielding potential. However, it is worthwhile to note that influenza glycoproteins are usually not as extensively glycosylated as on some other viral proteins, such as the HIV envelope protein or the SARS-CoV-2 S protein (15, 17, 103–105). The exact H5N1 construct prepared here contains a glycosylation site at N146. This is part of the 150 loop that borders the primary binding site (**Figure 1**). The representation in **Figure 2** shows that the glycans present at site N146 on each monomer have the combined capability to cover both NA binding sites, potentially thwarting the binding of small molecules. The results shown here display a moderate inhibitory effect due to glycans, but this effect would likely not be present in proteins whose glycans only reside far from the ligand binding sites, i.e. if the setup in **Figure 1** only contained the glycans at site 1 and site 3 on the bottom of the NA head. When examining the effect of glycan conformation on binding inhibition, the Glyprot glycans display a fairly vertical conformation. On the other hand, the glycans from the MD simulations bend backwards, away from the primary binding site on their own monomer and towards the secondary binding site of the adjacent monomer, as shown in **Figure 4**. Interestingly, this bend appears to be enough to inhibit primary site binding.

It has been previously shown that specific chemical modifications on the glycans can significantly change their flexibility (106–108). It has also been hypothesized that glycan flexibility plays a role in protein-receptor binding equilibria (87). Considering the scale of biological interactions that glycans participate in, it is likely that they would exploit their flexibility to facilitate these interactions. However, the glycan environment, and nearby steric clashes would conceivably affect this flexibility as well, introducing competing effects. Revisiting the input NA structure in **Figure 1**, we hypothesized that the glycan on top of each NA monomer (the oligomannose type glycans linked to site #1) would achieve a higher degree of flexibility than the two on the bottom of each monomer (the complex and hybrid type glycans linked to sites #2 and #3, respectively). Our reasoning was that these two may find steric restrictions on their flexibility, and that the placement on the glycan on the NA head would be more important than the type of glycan examined. Our results show this is not quite the case. The clusters in **Figure 3**, **Figure S6**, **Figure S7**, and **Figure S8**, show that, similar to the complex-type glycans (A-D2), the oligomannose-type glycans (A-D1) were quite flexible even though they were situated near the hybrid-type glycans (A-D3) on the bottom of the NA surface; this large degree of conformational freedom is backed up by previous work specifying that this flexibility is driven by the mannose(4)-α(1-3)-mannose(3) and the mannose(5)-α(1-6)-mannose(3) linkages (109). These are the linkages connecting the chitobiose glycan “stalk” to the two glycan “branches”. Finally, the hybrid-type glycans showed noticeably less conformational flexibility than either the oligomannose-type glycans or the complex-type glycans. Overall, the type of glycan and its specific linkages seemed to govern its flexibility more than potential nearby steric clashes. This agrees with previous work showing that unless there is a direct steric clash, inter-residue hydrogen bonds may have a larger effect governing glycan conformations (106, 109).

### Glycan effects on association rates

The results shown in **Figure 5** are consistent with diffusion controlled reactions, and show relatively high association rates. The space explored is consistent with the random walk nature of diffusion. The randomness of the ligand trajectories (from Brownian motion) and the small sizes of the ligands considered here minimize the effects of the glycans on binding. The rates for each ligand are mostly of similar orders of magnitude, with or without glycosylation. However, the glycan structures from the MD simulations show a moderate inhibition compared to the unglycosylated NA structure and the NA structure with glycan structures taken directly from the Glyprot web server. The extent of this inhibition ranges from a factor of about two to eight.

In general, glycans can decrease binding activity of viral proteins (3, 42, 44, 110). Due to their bulk and proximity to the primary ligand binding site, we hypothesized that, irrespective of conformation, the presence of glycans, particularly those near the binding sites, could substantially reduce ligand binding and removing these glycans would restore binding. What we found was a more nuanced picture. The NA constructs with glycan conformations from the Glyprot server showed similar binding rates to unglycosylated constructs. However, more realistic glycan conformations, extracted from the MD simulations, showed a moderate but noticeable decrease in association rate, k_on_, on the order of two to eight times. One may naturally question whether glycans would have the same effect on dissociation rate, k_off_. One previous study testing antibody binding to cancer cells showed that antibody binding was relatively insensitive to the presence of glycans, indicating a similar dampening of k_on_ and k_off_ due to the presence of glycans (9). In this study mentioned, the overall equilibrium constant K_D_ changed by less than a factor of two irrespective of the presence or absence of glycans (9). However, a different study done in the influenza membrane protein hemagglutinin showed that trimming the glycans from a standard length seen in HEK293 cells to a single monosaccharide decreases the equilibrium constant K_D_ by a factor of two to 48, depending on the receptor mimic used (42). This meant that the k_on_ and the k_off_ were not affected in the same way by the presence of glycans (42). Glycans are present in the antibody binding sites of both of the studies mentioned above; this is in contrast to our system where glycans are situated near the catalytic sites, but not directly inside them. With this in mind, it seems likely that the slight slowing of binding small ligands by the glycans would be similarly reflected in a slight slowing of release, so that the equilibrium constants for binding these molecules are relatively insensitive to the presence of glycans. In effect, this is because the presence of glycans near the binding sites should not change the ΔG in accordance with the Gibbs relationship. We hypothesize that our observed decrease in association rate is due to the glycans at glycosylation site N146 (site #2) as only those glycans are capable of sterically inhibiting the binding sites (**Figure 2**), and we assume the glycans at sites N88 (site #1) and N234 (site #3) do not impair binding. Taking the inhibition results discussed here with a different binding study using larger ligands for influenza NA, there appears to be a size dependence on this inhibitory potential: smaller ligands are not as affected as larger ligands (43). The key points here are that small molecules are not seriously impeded from binding by the glycans; future drug discovery efforts can be focused on the development of strong binders with correspondingly long lifetimes of binding. Modeling studies focused on small inhibitors are likely to be helpful, even when glycans are not included.

The results seen in **Figure 5** highlight the importance of using biologically-relevant glycan conformations relaxed on the protein structure as opposed to simply generating a glycan conformation and attaching it to the protein. Though this study did use static structures as per the BD setup, we would expect similar trends if this study were repeated using a dynamic MD environment since our BD trajectories already used the most highly-accessed glycan conformations gleaned from extensive MD sampling. Moreover, a study using mixed BD-MD simulations analyzing the association of oseltamivir and zanamivir to NA actually showed a less accurate k_on_ rate than our coarser study using only BD (54). We can rationalize that the slower binding kinetics seen in our systems with biologically-relevant glycan conformations (**Figure 5**) are due to the ligands having to maneuver around the glycans, even after running into them, and then continuing with the trajectory until reaching the binding site. This type of maneuverability can be seen in **Figure 6**.

**Figure 6.**
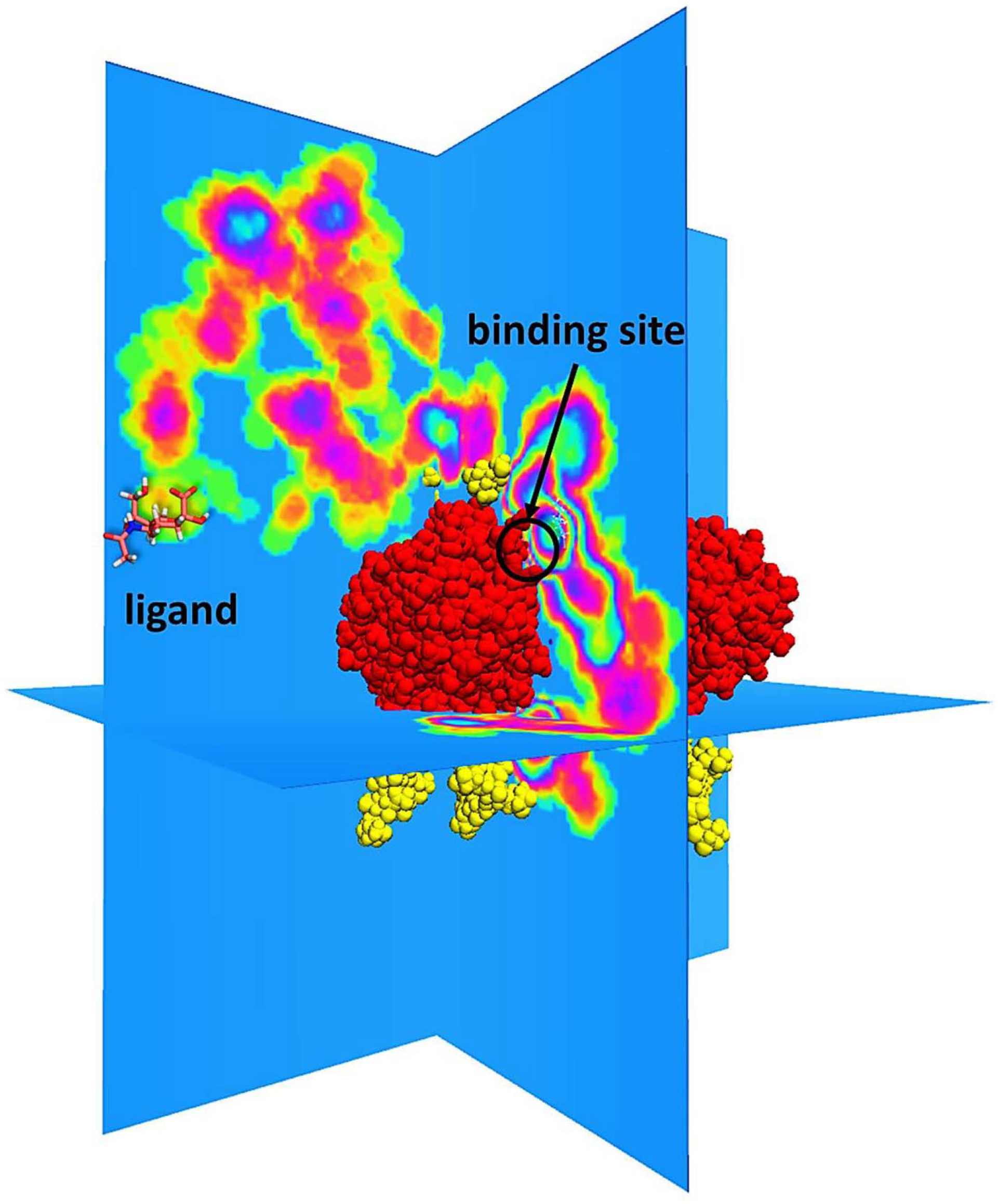
A sample successful trajectory, showing that the ligand explores significant space before finding the binding site. This sample NA contains glycans from the Glyprot server. Sialic acid (not to scale) is superimposed on the left side of the image for clarity. Because of the space explored, it makes sense that a blockage in part of the path won’t significantly affect the association rate. Here, the protein is in red and the glycans are in yellow. The blue planes are used for creating subsections of 3D space into 2D space. The colors on the planes indicate how often the ligand has spent in that 2D space, with the lime green inside magenta circles being the most occupied.

### Ligand binding at the primary and secondary sites

We generated BD trajectories that could end with the ligand binding to the primary site (**Figure 5A**), the secondary site (**Figure 5B**), or either site (**Figure 5C**) on any monomer. Using this setup, we were able to differentiate binding between the primary and secondary sites, and in fact found an additive binding mode when examining both sites concurrently. By simply adding up the association rates observed for the primary site (**Figure 5A**) to the analogous simulation run to the secondary site (**Figure 5B**), the association rate to both sites (**Figure 5C**) can be roughly obtained. We do not see any evidence of a further increase in association rate using both sites, showing that the presence of a proximal binding site does not influence association rate, either for the primary site or the secondary site.

Our primary site binding results show two conclusions supported by literature. In **Figure 5A** we see that oseltamivir associates faster than zanamivir, as has been seen in experimental kinetics studies (60). Moreover, we see faster binding of oseltamivir than sialic acid. This is qualitatively in agreement with an NMR study showing that oseltamivir outcompetes α(2,3)-sialyllactose in binding to the avian NA active site (97). It is not immediately clear which atoms on the ligands drive their binding differences.

Ligand binding to the secondary site has not been extensively studied, but it does not appear to have catalytic activity (111, 112). Focusing on the secondary site, our results show three important findings. We first see that binding to the secondary site is slower than to the primary site, if binding is seen at all (**Figure 5**). We do not see secondary site binding for oseltamivir and very little for zanamivir, though this may be as they are at the lower detection limit of our method. Furthermore, we see that sialic acid binds faster to the secondary site than oseltamivir, which is in agreement with one study showing that α(2,3)-sialyllactose outcompetes oseltamivir for binding to the avian NA secondary site (97). A more recent study goes further and does not show any binding of oseltamivir to the avian NA secondary site (98). However, we caution that a small amount of drug binding, likely only with zanamivir, may occur to the secondary site, as seen with zanamivir bound in the secondary site in the unpublished crystal structure PDB 2CML, and also seen in **Figure 5B**. Secondly, in the small amount of secondary site binding seen (**Figure 5B**), glycans are actually capable of enhancing or inhibiting binding, foreshadowing the complex role glycans play in ligand binding. Finally, there appears to be a small conformational dependence on association rate, but this is only seen towards the secondary site (**Figure 5B**). We used two different conformations of sialic acid for these binding studies. The boat conformation was crystallized in the active site and the chair conformation was crystallized in the secondary site. In our results we see the sialic acid chair conformation actually shows fractionally higher binding to the primary site than the boat conformation (**Figure 5A**). Conversely, only the boat conformation shows binding to the secondary site; the chair conformation does not register binding at all (**Figure 5B**). However, we caution that these results may be because sialic acid was crystallized in a different strain of avian NA than we used in our studies. Taken together, these results show that the exact ligand conformation upon approach to the binding site may not match the crystallized binding pose, but the results we present here do not permit us to explore this note or further explain a conformational dependence on binding.

Comparing the association rates in **Figure 5** one may naturally query the competition in association rates between the primary and secondary sites. We see faster association to the primary site than the secondary site, which is not in agreement with two previous BD simulation studies (52, 53). However, the methodology of our study differs from these two studies, and from this we can unify the difference. Those BD studies showed that ligands reach a distance of 7.5 Å away from the secondary site faster than to the primary site. We then show that ligands reach a distance of 3.228 Å away from the primary site faster than the secondary site, though we would like to note that our paper and the Sung et al. paper assigned charges for the BD trajectories according to the AMBER99 force field and the Amaro et al. paper assigned charges according to the CHARMM36 force field (52, 53). Taken together, the secondary site appears to contain stronger long-range electrostatics to draw in ligands, but when the ligands approach the binding sites and sterics come into play, it appears to be more favorable for ligands to move closer to the primary site than the secondary site, assuming the rigid body approximations applied herein.

Considering the fact that the realistic substrates NA encounters will exhibit multivalent binding, one previous study showed that the secondary site improved avian NA enzymatic activity in removing sialic acid both from soluble macromolecular substrates and from cells (113). Another study confirmed that the binding in the secondary site improved catalytic activity against multivalent substrates (114). Other previous studies have suggested that the secondary site enhances the overall NA catalytic activity by binding substrates and bringing them close to the catalytic primary site (111, 113–116). Taking the studies above with our results, we postulate that multivalent cleavage will occur in a stepwise manner (**Figure 7**). The first association event of the multivalent substrate, such as sialylated cell surface receptors, will bind to the primary site, and then to the secondary site. After sialidase cleavage occurs in the primary site, the cleaved glycan branch will dissociate. Then the sialylated glycan branch bound in the secondary site will be transferred to the primary site, as suggested previously (111, 113–115). After this passage, cleavage will again occur, and the full glycan will be released, finishing the enzymatic cycle. This mechanism is in disagreement with a previously proposed mechanism, which postulates that both binding sites will not be bound simultaneously (115). However, we feel there is a greater body of literature suggesting that binding both sites simultaneously increases catalytic activity. We note that our proposed binding mechanism may be muddied in the case of multivalent ligands with viral glycans situated near the binding sites; in this case, the glycans may sterically inhibit multivalent binding, slowing down enzymatic activity and attenuating the replication cycle. In the case of monovalent binders, such as the inhibitors oseltamivir and zanamivir, we show in **Figure 5** that association will happen to the primary site faster than to the secondary site. This appears to be biologically viable considering that previous studies have showed that the secondary site activity has no effect on enzymatic activity for monovalent substrates (98, 117–119). As the primary site is the main site of enzymatic activity, it is reasonable to assume that ligands would preferentially bind to the primary site over the secondary site; reducing transfers of ligands between the binding sites would ostensibly increase catalytic activity and efficiency. Taken together, abolishing the secondary site in avian NA will not affect monovalent substrates such as influenza drugs as these associate faster to the primary site anyways, which our results confirm. To exposit this a different way, influenza drugs will preferentially block primary site binding over secondary site binding.

**Figure 7.**
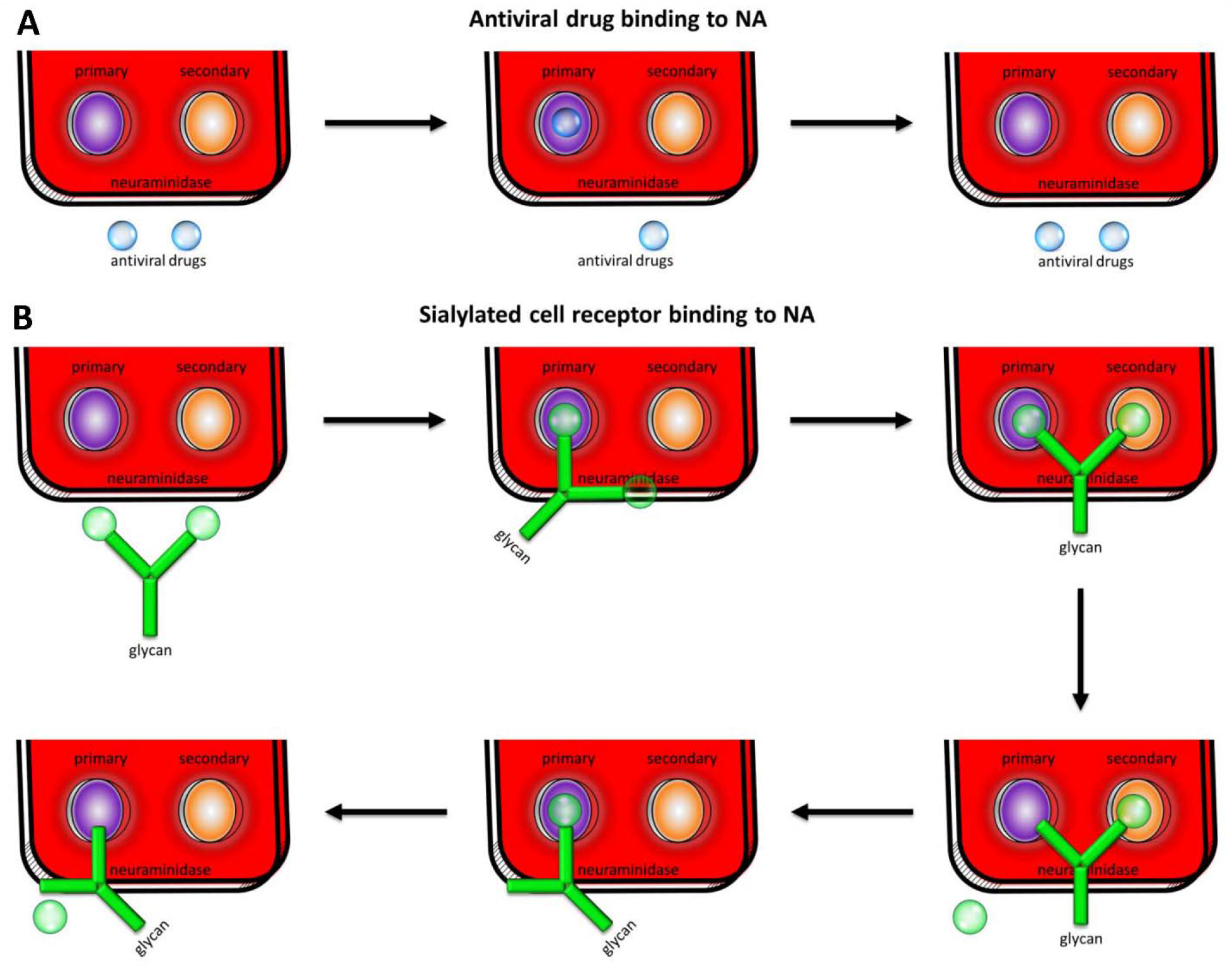
Ligand binding mechanisms to neuraminidase for monovalent binders (A) and multivalent binders (B). A free monovalent binder will associate to the primary site over the secondary site (A). This monovalent binder will release from the primary site before a second monovalent binder will associate to the secondary site. Glycans, with their sialic acid tips, are an example of a multivalent binder (B). Similar to the monovalent binders, the first multivalent binding event will occur to the primary site. Next, the second sialic acid tip binds to the secondary site. With both sites bound, the sialic acid in the primary site is cleaved and released. The sialic acid bound in the secondary site is then transferred to the primary site. Finally, the second sialic acid is cleaved and released, and the enzymatic cycle is complete.

## Conclusion

In this work, we created NA systems with varying glycan conformations, and also without the presence of glycans. These glycans are capable of covering much of the surface area of NA. Their conformational flexibility is dependent on their glycan type, not necessarily their spatial position. The glycosylated systems showed moderate inhibition of ligands to the primary binding site. Finally, we propose a new binding mechanism for multivalent binders to NA, such as cell surface receptors. These results have implications for future drug development, the overall understanding of glycans, and the NA enzymatic mechanism. Much sustained effort has gone into developing NA inhibitors, and will continue to do so in the future. Measuring the binding of a potential drug is an important step in the drug discovery process. However, most drug discovery efforts have not taken into account viral glycans. Neglecting this effect can lead to a surprising drop in drug binding (9). Our work shows that glycans can have an inhibitory effect on influenza NA primary site binding. There have already been a number of studies using multivalent binders as NA antivirals (120–125). With the results shown here, we recommend future work on multivalent NA drugs, to focus on developing strong binders with a long lifetime, regardless of the presence or absence of glycans. With the detection limitations of our study, we cannot conclude how glycans affect secondary site binding, although we believe binding to the secondary site will be slower than binding to the primary site (**Figure 5**). However, it follows from these results that glycans could evoke a secondary site binding inhibition similar to the primary site. In summary, this work examines glycan inhibition on drug binding, compares the drug binding interplay between two binding sites, and proposes a new mechanism of ligand binding to NA.

## Author Contributions

C.S., R.E.A., and J.A.M. designed the experiments and performed analysis. C.S. ran the simulations and wrote the manuscript with contributions from all authors. L.C., R.K., and G.H. provided critical support necessary for the simulations to run. All authors approved the final version of the manuscript.

## Acknowledgements

C.S. would like to thank Zied Gaieb for valuable discussions. This work was supported in part by grants from the National Institutes of Health, USA (NIH grant T32EB009380 to C.S. and GM031749 to J.A.M. and G.H.). This material is based upon work supported by the National Science Foundation Graduate Research Fellowship Program under Grant No. DGE-1650112 to C.S. Any opinions, findings, and conclusions or recommendations expressed in this material are those of the author(s) and do not necessarily reflect the views of the National Science Foundation. This work used the Extreme Science and Engineering Discovery Environment (XSEDE), which is supported by National Science Foundation grant number ACI-1548562. This work used the Extreme Science and Engineering Discovery Environment (XSEDE) Comet at the San Diego Supercomputer Center (SDSC) through allocation csd373 to R.E.A. R.E.A. has equity interest in and is a cofounder of and on the scientific advisory board of Actavalon.

