## Supplementary Material for "Multiscale simulations examining glycan shield effects on drug binding to influenza neuraminidase"

Co-corresponding author: Christian Seitz

Twitter handle: @chem_christian

Co-corresponding author: Rommie E. Amaro

Twitter handle: @RommieAmaro

**Supporting Material**

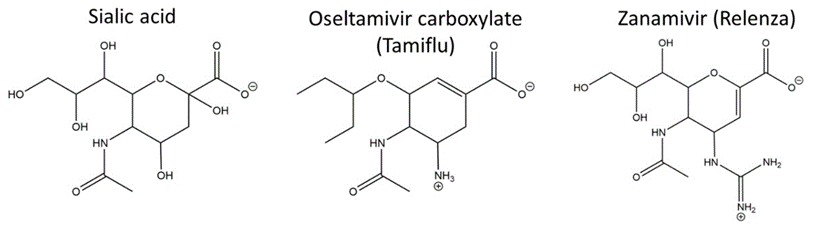

**Figure S1**. **2D structures of the ligands used in this study.**

**Table S1**. **Ions used in the implicit solvent in the BD trajectories.**

| **Ion** | **Charge** | **Concentration** | **Radius** |
| --- | --- | --- | --- |
| Ca^2+^ | 2.000 | 0.005 M | 1.7131 Å |
| Cl^-^ | -1.000 | 0.010 M | 1.7350 Å |
| Na^+^ | 1.000 | 0.024 M | 1.8680 Å |
| MES^-^ | -1.000 | 0.024 M | 4.0070 Å |

**Table S2. Grid spacing for each system**. The grid spacing is defined as the fine grid length divided by the number of grid points.

| **System** | **Grid spacing** |
| --- | --- |
| Unglycosylated NA | 0.61 x 0.61 x 0.66 Å |
| NA with Glyprot glycans | 0.80 x 0.80 x 0.83 Å |
| NA with MD conf1 glycans | 0.59 x 0.59 x 0.68 Å |
| NA with MD conf2 glycans | 0.60 x 0.61 x 0.70 Å |
| NA with MD conf3 glycans | 0.60 x 0.61 x 0.70 Å |

**Table S3. b radius and q radius values for each system**. The distance is the distance away from the hydrodynamic center of the protein.

| **System** | **b radius** | **q radius** |
| --- | --- | --- |
| Unglycosylated NA | 109.047 Å | 119.952 Å |
| NA with Glyprot glycans | 109.310 Å | 120.241 Å |
| NA with MD conf1 glycans | 111.999 Å | 123.199 Å |
| NA with MD conf2 glycans | 110.382 Å | 121.420 Å |
| NA with MD conf3 glycans | 110.098 Å | 121.108 Å |

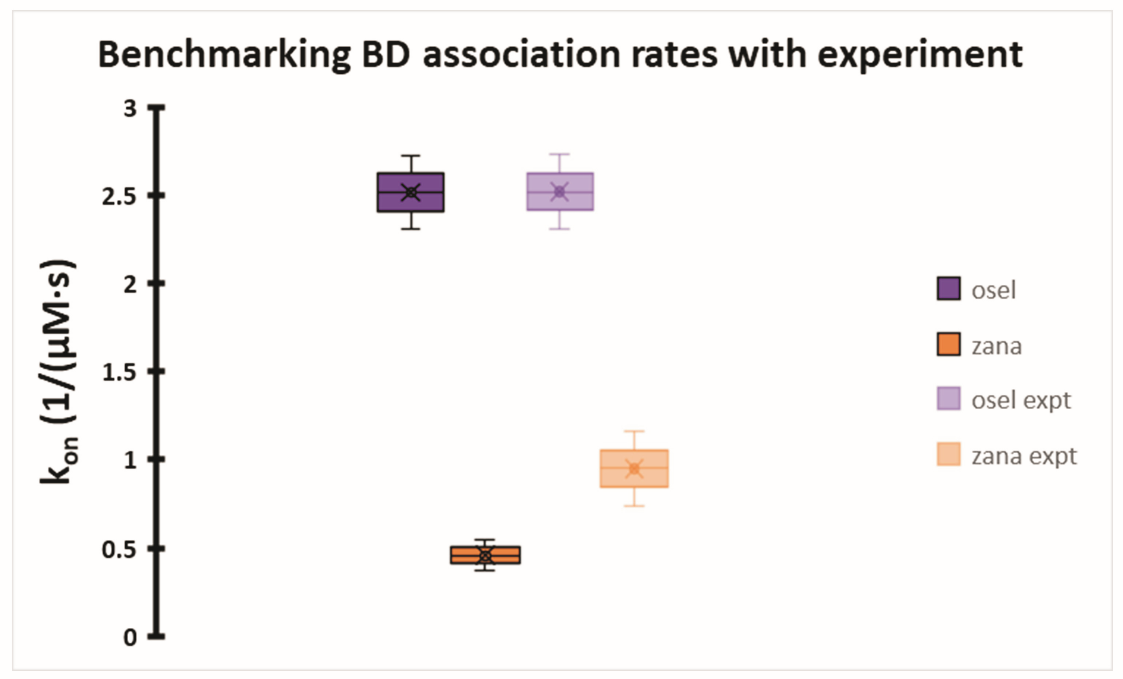

**Figure S2**. **Comparison of our computed association rates with previously published experimental association rates**. The computed oseltamivir rate (purple) is within the level of error of the experimental oseltamivir rate (faded purple). The computed zanamivir rate (orange) is similar to the experimental zanamivir rate (faded orange).

Overall, the primary site binding parameters were transferred from oseltamivir to zanamivir, correcting for the replacement of oseltamivir’s isopentanetriol ring substituent with zanamivir’s 1,2,3-pentanetriol ring substituent, as seen in **Figure S1**. They were then transferred to sialic acid, accounting for the methanetriamine ring substituent (**Figure S1**). The secondary site binding parameters were determined for sialic acid before being transferred to oseltamivir and zanamivir. These choices were made due to the availability of crystal structures.

The primary binding site contacts for oseltamivir are presented in **Table S3**. The oseltamivir isopentanetriol group is replaced by the sialic acid 08 and 09 oxygen atoms in binding pairs for both the primary and secondary sites (**Table S5** and **Table S8**). This isopentanetriol group in oseltamivir appears to form hydrophobic contacts with E276 of the neuraminidase instead of the hydrogen bonding that zanamivir and sialic acid shows, so these hydrophobic contacts were approximated as having contacts to the C91, C8 and C82 atoms as shown above. The zanamivir (**Table S4**) binding pairs are based off of those for oseltamivir.

The 1MWE crystal structure shows the tryptophan’s conjugated benzene ring having an interaction with the C11 atom of sialic acid in the secondary binding site. We approximated this putative hydrophobic interaction by picking three evenly spaced carbon atoms on the tryptophan ring as shown in **Table S8**. These carbons were picked to be away from the indole ring, as they would be less polar. The secondary site contacts in the 1MWE crystal structure also appear to show hydrophobic contacts between NA residue S370 and the ligand. These were approximated by selecting the α-carbon of S370, as it would be less polar than the side chain carbon. Lastly, the secondary site 1MWE crystal structure showed hydrophobic interactions between K432 and sialic acid. These were approximated by selecting the CG atom of K432, which is in the middle of the side chain away from the nitrogen. The binding pairs for oseltamivir (**Table S5**) and zanamivir (**Table S7**) are based off those for sialic acid.

In summary, the primary and secondary site binding pairs for these ligands are set to be analogs of each other. The NA binding pair atoms were kept constant across each ligand for consistency.

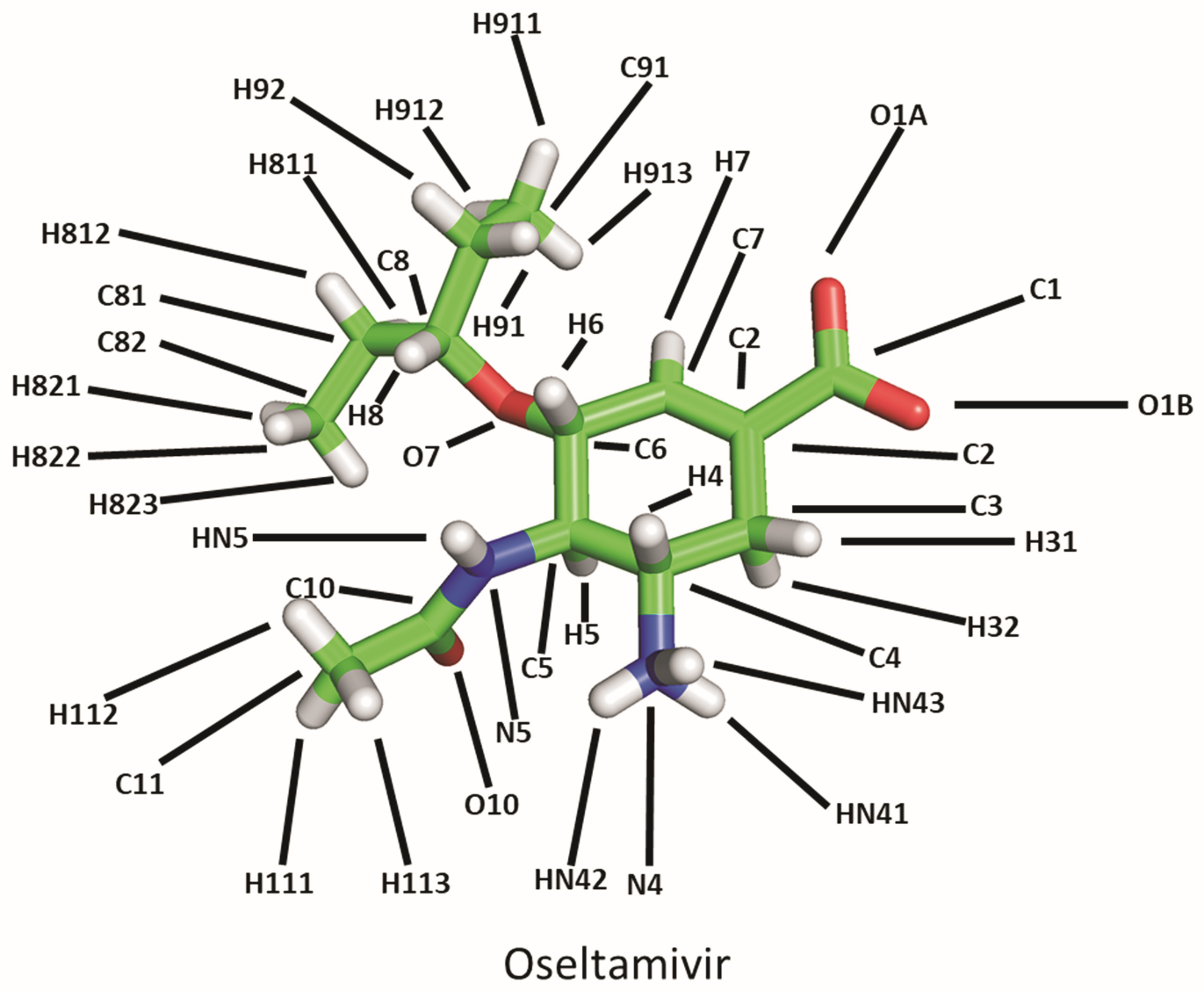

**Figure S3**. **Structure of oseltamivir**.

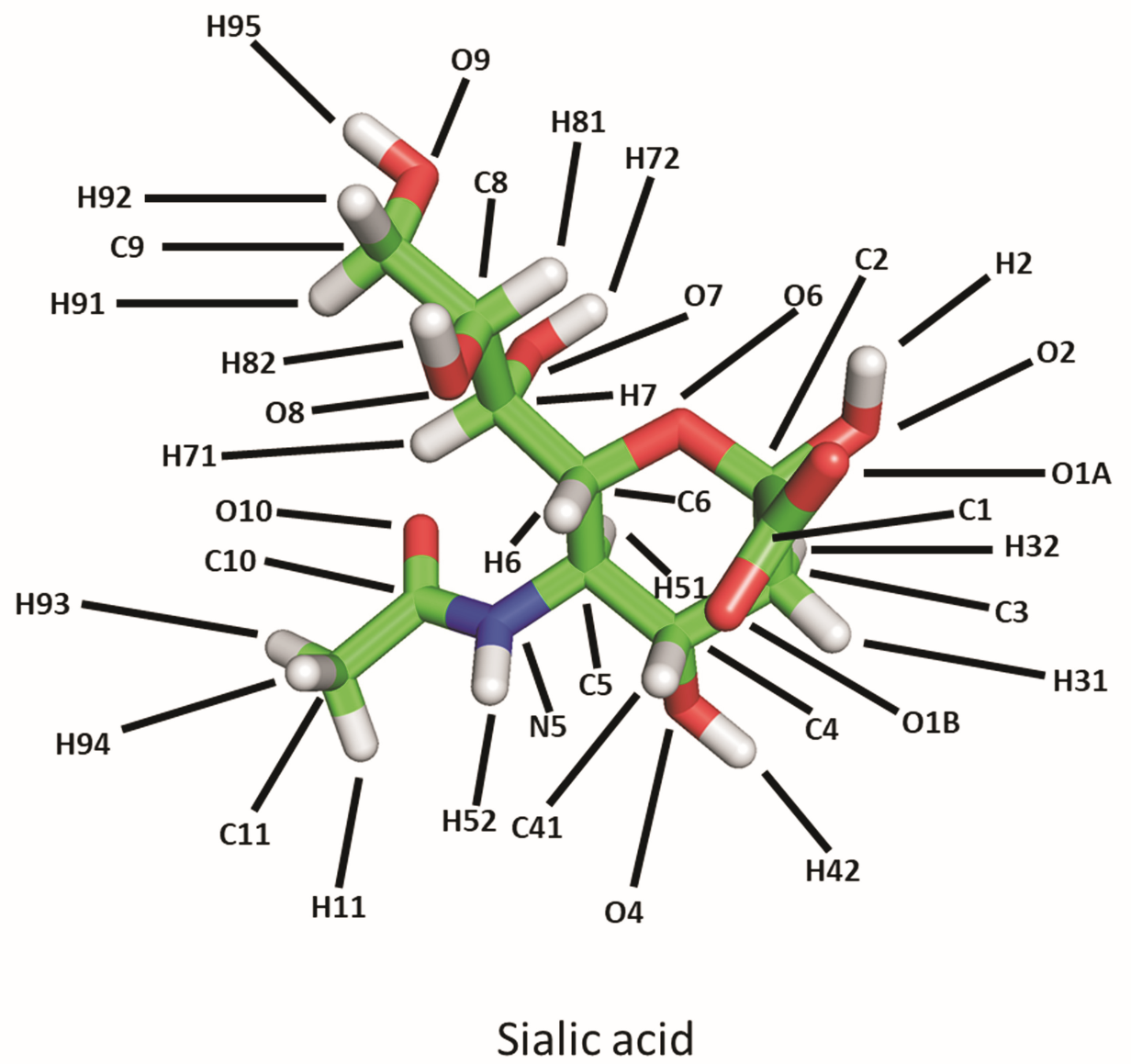

**Figure S4**. **Structure of sialic acid**.

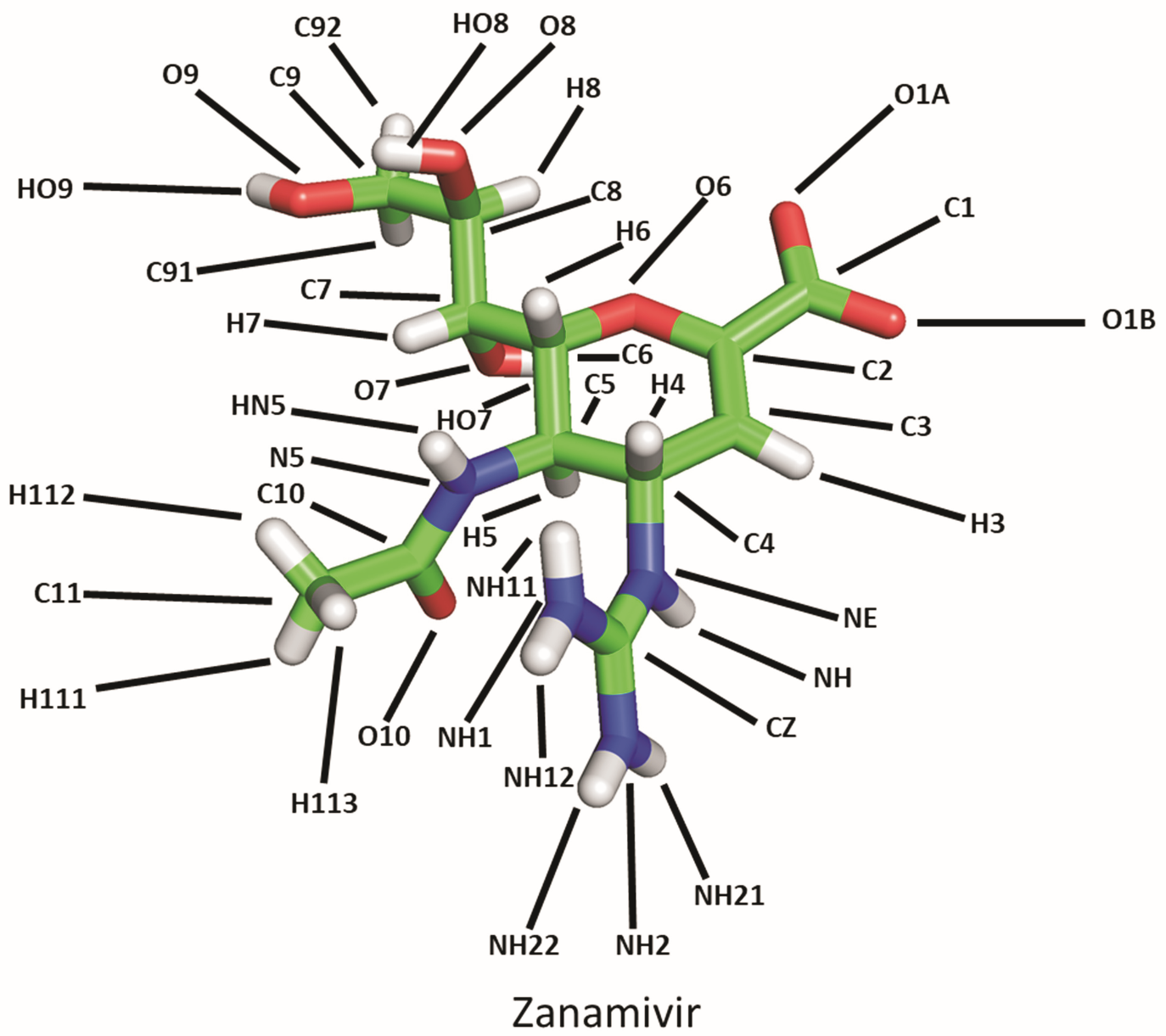

**Figure S5**. **Structure of zanamivir**.

**Table S4. Primary site atom pairs for oseltamivir**.

| **Neuraminidase residue** | **Residue atom** | **Oseltamivir atom** |
| --- | --- | --- |
| R118 | Nη_1_ | O1A |
| R118 | Nη_1_ | O1B |
| R118 | Nη_2_ | O1A |
| R118 | Nη_2_ | O1B |
| R152 | Nη_1_ | O10 |
| R152 | Nη_2_ | O10 |
| E276 | Cβ | C8 |
| E276 | Cβ | C82 |
| E276 | Cβ | C91 |
| R292 | Nη_1_ | O1A |
| R292 | Nη_1_ | O1B |
| R292 | Nη_2_ | O1A |
| R292 | Nη_2_ | O1B |
| Y347 | Oη | O1A |
| Y347 | Oη | O1B |
| R371 | Nη_1_ | O1A |
| R371 | Nη_1_ | O1B |
| R371 | Nη_2_ | O1A |
| R371 | Nη_2_ | O1B |
| R371 | Nε | O1B |
| R371 | Nε | O1A |
| Y406 | Oη | O1A |
| Y406 | Oη | O1B |

**Table S5**. **Primary site atom pairs for sialic acid**.

| **Neuraminidase residue** | **Residue atom** | **Sialic acid atom** |
| --- | --- | --- |
| R118 | Nη_1_ | O1A |
| R118 | Nη_1_ | O1B |
| R118 | Nη_2_ | O1A |
| R118 | Nη_2_ | O1B |
| R152 | Nη_1_ | O10 |
| R152 | Nη_2_ | O10 |
| E276 | Oε_1_ | O8 |
| E276 | Oε_2_ | O8 |
| E276 | Oε_1_ | O9 |
| E276 | Oε_2_ | O9 |
| R292 | Nη_1_ | O1A |
| R292 | Nη_1_ | O1B |
| R292 | Nη_2_ | O1A |
| R292 | Nη_2_ | O1B |
| Y347 | Oη | O1A |
| Y347 | Oη | O1B |
| R371 | Nη_1_ | O1A |
| R371 | Nη_1_ | O1B |
| R371 | Nη_2_ | O1A |
| R371 | Nη_2_ | O1B |
| R371 | Nε | O1B |
| R371 | Nε | O1A |
| Y406 | Oη | O1A |
| Y406 | Oη | O1B |

**Table S6**. **Primary site atom pairs for zanamivir**.

| **Neuraminidase residue** | **Residue atom** | **Zanamivir atom** |
| --- | --- | --- |
| R118 | Nη_1_ | O1A |
| R118 | Nη_1_ | O1B |
| R118 | Nη_2_ | O1A |
| R118 | Nη_2_ | O1B |
| R152 | Nη_1_ | O10 |
| R152 | Nη_2_ | O10 |
| E276 | Oε_1_ | O8 |
| E276 | Oε_2_ | O8 |
| E276 | Oε_1_ | O9 |
| E276 | Oε_2_ | O9 |
| R292 | Nη_1_ | O1A |
| R292 | Nη_1_ | O1B |
| R292 | Nη_2_ | O1A |
| R292 | Nη_2_ | O1B |
| Y347 | Oη | O1A |
| Y347 | Oη | O1B |
| R371 | Nη_1_ | O1A |
| R371 | Nη_1_ | O1B |
| R371 | Nη_2_ | O1A |
| R371 | Nη_2_ | O1B |
| R371 | Nε | O1B |
| R371 | Nε | O1A |
| Y406 | Oη | O1A |
| Y406 | Oη | O1B |

**Table S7**. **Secondary site atom pairs for sialic acid**.

| **Neuraminidase residue** | **Residue atom** | **Sialic acid atom** |
| --- | --- | --- |
| S367 | OG | O1A |
| S367 | OG | O1B |
| S370 | CA | C91 |
| S370 | CA | C8 |
| S370 | CA | C82 |
| S372 | OG | N5 |
| W403 | CD2 | C11 |
| W403 | CZ2 | C11 |
| W403 | CZ3 | C11 |
| K432 | CG | O8 |
| K432 | CG | O9 |

**Table S8. Secondary site atom pairs for oseltamivir**.

| **Neuraminidase residue** | **Residue atom** | **Oseltamivir atom** |
| --- | --- | --- |
| S367 | OG | O1A |
| S367 | OG | O1B |
| S370 | CA | C91 |
| S370 | CA | C8 |
| S370 | CA | C82 |
| S372 | OG | N5 |
| W403 | CD2 | C11 |
| W403 | CZ2 | C11 |
| W403 | CZ3 | C11 |
| K432 | CG | C91 |
| K432 | CG | C8 |
| K432 | CG | C82 |

**Table S9**. **Secondary site atom pairs for zanamivir**.

| **Neuraminidase residue** | **Residue atom** | **Zanamivir atom** |
| --- | --- | --- |
| S367 | OG | O1A |
| S367 | OG | O1B |
| S370 | CA | C91 |
| S370 | CA | C8 |
| S370 | CA | C82 |
| S372 | OG | N5 |
| W403 | CD2 | C11 |
| W403 | CZ2 | C11 |
| W403 | CZ3 | C11 |
| K432 | CG | O8 |
| K432 | CG | O9 |

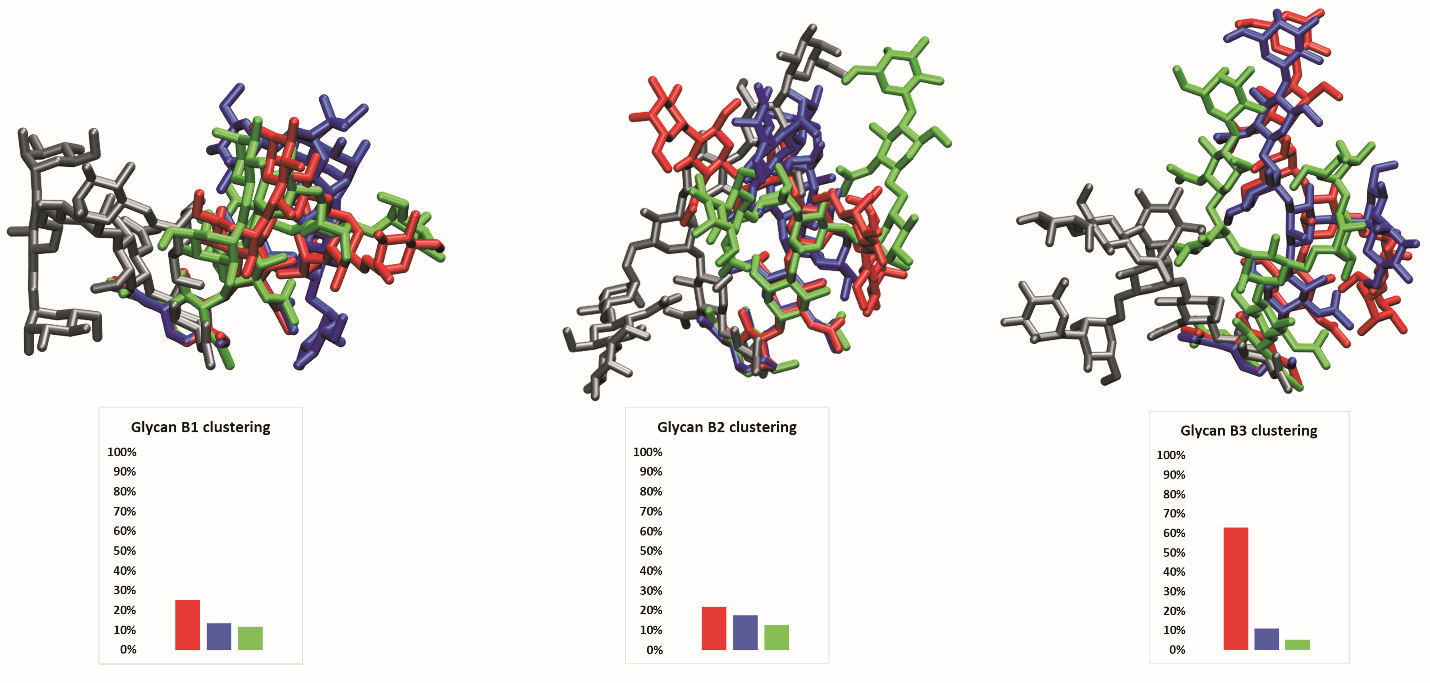

**Figure S6**. **Clustered glycans on monomer B**. The Glyprot structure is in gray, while the other colors represent clusters from the MD simulations.

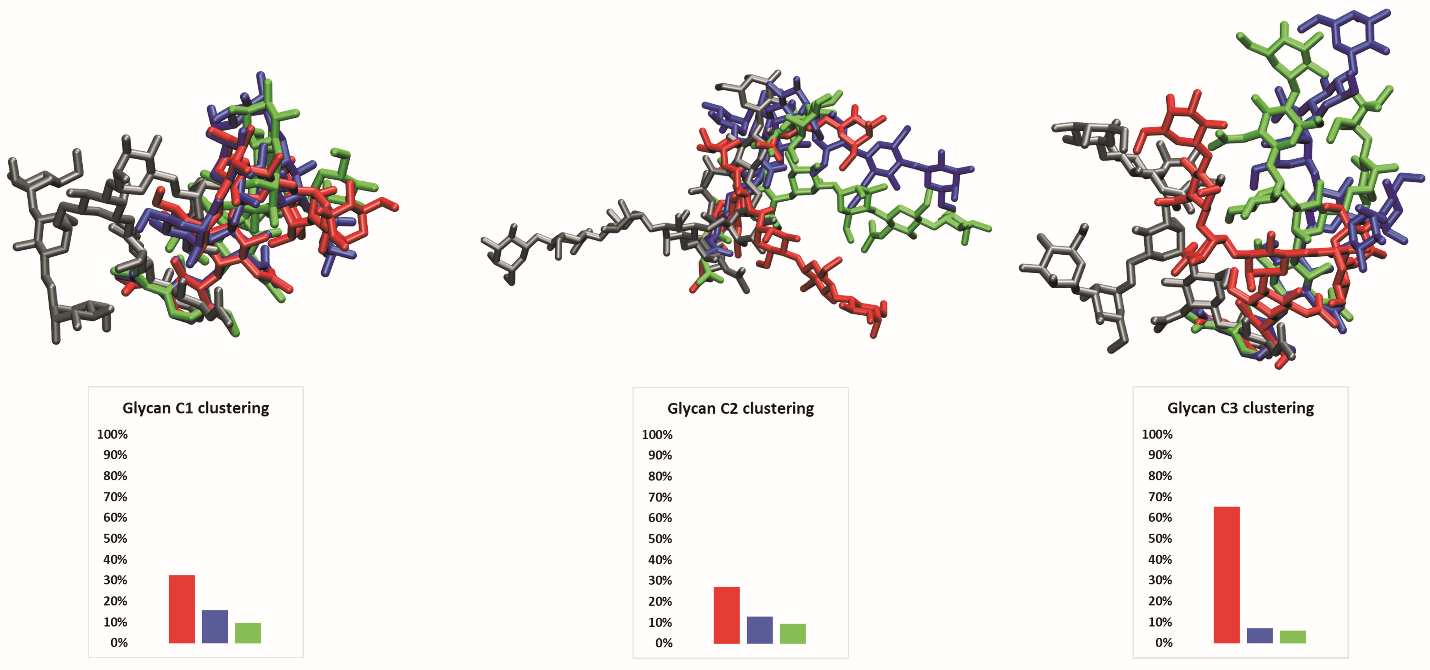

**Figure S7**. **Clustered glycans on monomer C**. The Glyprot structure is in gray, while the other colors represent clusters from the MD simulations.

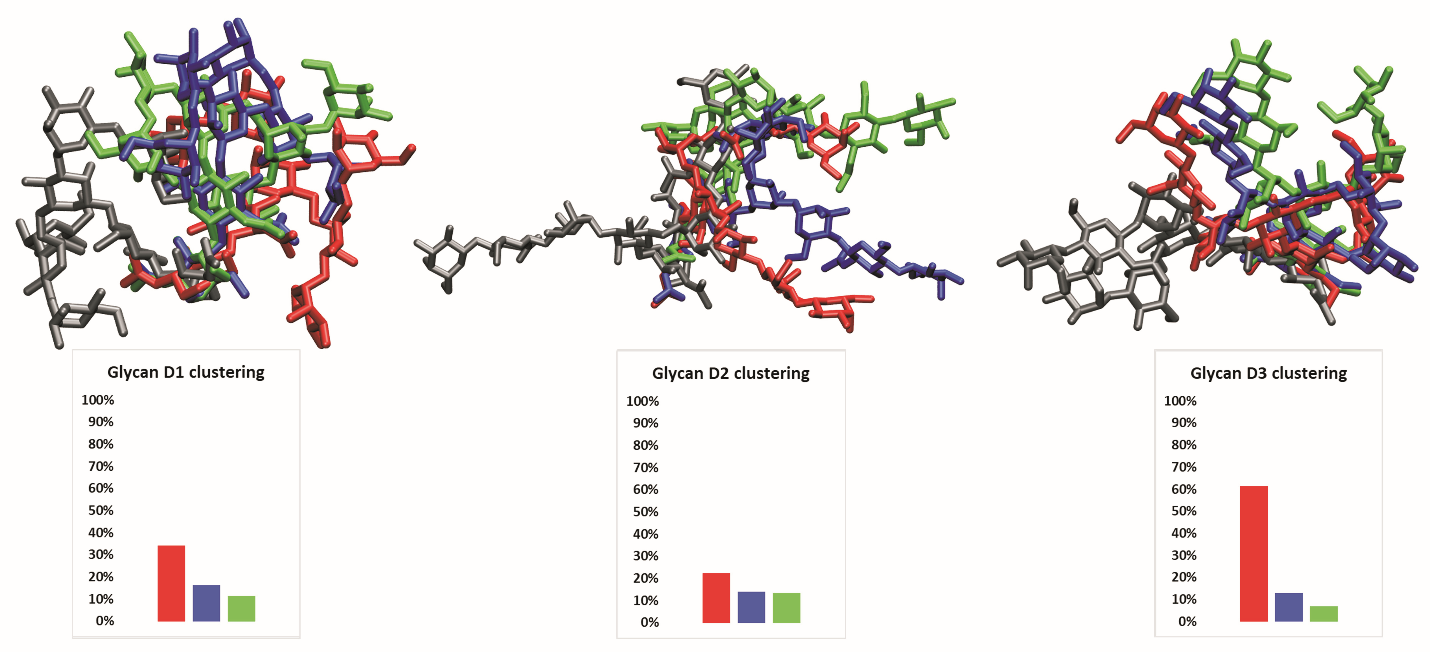

**Figure S8**. **Clustered glycans on monomer D**. The Glyprot structure is in gray, while the other colors represent clusters from the MD simulations.

**Table S10**. **Association rate data for oseltamivir.** The files containing this data are provided.

| **System** | **k_on_** | | | **Probability** | | | **Total binding events** |
| --- | --- | --- | --- | --- | --- | --- | --- |
|  | **low** | **mean** | **high** | **low** | **mean** | **high** |  |
| **Primary site** | | | | | | | |
| Unglycosylated NA | 3.66/µM∙s | 3.93/µM∙s | 4.19/µM∙s | 8.28 x 10^-5^ | 8.88 x 10^-5^ | 9.48 x 10^-5^ | 888 |
| NA with Glyprot glycans | 2.31/µM∙s | 2.52/µM∙s | 2.73/µM∙s | 5.24 x 10^-5^ | 5.72 x 10^-5^ | 6.20 x 10^-5^ | 572 |
| NA with MD conf1 glycans | 1.22/µM∙s | 1.38/µM∙s | 1.53/µM∙s | 2.68 x 10^-5^ | 3.03 x 10^-5^ | 3.38 x 10^-5^ | 303 |
| NA with MD conf2 glycans | 0.47/µM∙s | 0.57/µM∙s | 0.68/µM∙s | 1.06 x 10^-5^ | 1.29 x 10^-5^ | 1.52 x 10^-5^ | 129 |
| NA with MD conf3 glycans | 1.39/µM∙s | 1.55/µM∙s | 1.72/µM∙s | 3.13 x 10^-5^ | 3.50 x 10^-5^ | 3.87 x 10^-5^ | 350 |
| **Secondary site** | | | | | | | |
| Unglycosylated NA | 0.00/µM∙s | 0.00/µM∙s | 0.00/µM∙s | 0.00 x 10^-5^ | 0.00 x 10^-5^ | 0.00 x 10^-5^ | 0 |
| NA with Glyprot glycans | 0.00/µM∙s | 0.00/µM∙s | 0.00/µM∙s | 0.00 x 10^-5^ | 0.00 x 10^-5^ | 0.00 x 10^-5^ | 0 |
| NA with MD conf1 glycans | 0.00/µM∙s | 0.00/µM∙s | 0.00/µM∙s | 0.00 x 10^-5^ | 0.00 x 10^-5^ | 0.00 x 10^-5^ | 0 |
| NA with MD conf2 glycans | 0.00/µM∙s | 0.00/µM∙s | 0.00/µM∙s | 0.00 x 10^-5^ | 0.00 x 10^-5^ | 0.00 x 10^-5^ | 0 |
| NA with MD conf3 glycans | 0.00/µM∙s | 0.00/µM∙s | 0.00/µM∙s | 0.00 x 10^-5^ | 0.00 x 10^-5^ | 0.00 x 10^-5^ | 0 |
| **Combined sites** | | | | | | | |
| Unglycosylated NA | 3.67/µM∙s | 3.93/µM∙s | 4.19/µM∙s | 8.29 x 10^-5^ | 8.89 x 10^-5^ | 9.49 x 10^-5^ | 889 |
| NA with Glyprot glycans | 2.49/µM∙s | 2.70/µM∙s | 2.92/µM∙s | 5.65 x 10^-5^ | 6.15 x 10^-5^ | 6.65 x 10^-5^ | 615 |
| NA with MD conf1 glycans | 1.47/µM∙s | 1.64/µM∙s | 1.82/µM∙s | 3.24 x 10^-5^ | 3.62 x 10^-5^ | 4.00 x 10^-5^ | 362 |
| NA with MD conf2 glycans | 0.48/µM∙s | 0.58/µM∙s | 0.68/µM∙s | 1.07 x 10^-5^ | 1.30 x 10^-5^ | 1.53 x 10^-5^ | 130 |
| NA with MD conf3 glycans | 1.42/µM∙s | 1.59/µM∙s | 1.75/µM∙s | 3.19 x 10^-5^ | 3.57 x 10^-5^ | 3.95 x 10^-5^ | 357 |

**Table S11**. **Association rate data for zanamivir**. The files containing this data are provided.

| **System** | **k_on_** | | | **Probability** | | | **Total binding events** |
| --- | --- | --- | --- | --- | --- | --- | --- |
|  | **low** | **mean** | **high** | **low** | **mean** | **high** |  |
| **Primary site** | | | | | | | |
| Unglycosylated NA | 0.28/µM∙s | 0.36/µM∙s | 0.44/µM∙s | 0.66 x 10^-5^ | 0.84 x 10^-5^ | 1.02 x 10^-5^ | 84 |
| NA with Glyprot glycans | 0.37/µM∙s | 0.46/µM∙s | 0.55/µM∙s | 0.85 x 10^-5^ | 1.06 x 10^-5^ | 1.27 x 10^-5^ | 106 |
| NA with MD conf1 glycans | 0.10/µM∙s | 0.16/µM∙s | 0.21/µM∙s | 0.23 x 10^-5^ | 0.35 x 10^-5^ | 0.47 x 10^-5^ | 35 |
| NA with MD conf2 glycans | 0.03/µM∙s | 0.07/µM∙s | 0.10/µM∙s | 0.07 x 10^-5^ | 0.15 x 10^-5^ | 0.23 x 10^-5^ | 15 |
| NA with MD conf3 glycans | 0.11/µM∙s | 0.17/µM∙s | 0.22/µM∙s | 0.26 x 10^-5^ | 0.38 x 10^-5^ | 0.50 x 10^-5^ | 38 |
| **Secondary site** | | | | | | | |
| Unglycosylated NA | 0.00/µM∙s | 0.00/µM∙s | 0.00/µM∙s | 0.00 x 10^-5^ | 0.00 x 10^-5^ | 0.00 x 10^-5^ | 0 |
| NA with Glyprot glycans | 0.00/µM∙s | 0.00/µM∙s | 0.00/µM∙s | 0.00 x 10^-5^ | 0.00 x 10^-5^ | 0.00 x 10^-5^ | 0 |
| NA with MD conf1 glycans | 0.01/µM∙s | 0.04/µM∙s | 0.07/µM∙s | 0.03 x 10^-5^ | 0.09 x 10^-5^ | 0.15 x 10^-5^ | 9 |
| NA with MD conf2 glycans | 0.00/µM∙s | 0.01/µM∙s | 0.02/µM∙s | -0.01 x 10^-5^ | 0.02 x 10^-5^ | 0.05 x 10^-5^ | 2 |
| NA with MD conf3 glycans | 0.02/µM∙s | 0.05/µM∙s | 0.08/µM∙s | 0.05x 10^-5^ | 0.12 x 10^-5^ | 0.19 x 10^-5^ | 12 |
| **Combined sites** | | | | | | | |
| Unglycosylated NA | 0.27/µM∙s | 0.35/µM∙s | 0.43/µM∙s | 0.63 x 10^-5^ | 0.81 x 10^-5^ | 0.99 x 10^-5^ | 81 |
| NA with Glyprot glycans | 0.27/µM∙s | 0.35/µM∙s | 0.43/µM∙s | 0.64 x 10^-5^ | 0.82 x 10^-5^ | 1.00 x 10^-5^ | 82 |
| NA with MD conf1 glycans | 0.16/µM∙s | 0.22/µM∙s | 0.29/µM∙s | 0.36 x 10^-5^ | 0.50 x 10^-5^ | 0.64 x 10^-5^ | 50 |
| NA with MD conf2 glycans | 0.03/µM∙s | 0.07/µM∙s | 0.10/µM∙s | 0.07 x 10^-5^ | 0.15 x 10^-5^ | 0.23 x 10^-5^ | 15 |
| NA with MD conf3 glycans | 0.17/µM∙s | 0.24/µM∙s | 0.30/µM∙s | 0.40 x 10^-5^ | 0.55 x 10^-5^ | 0.70 x 10^-5^ | 55 |

**Table S12**. **Association rate data for sialic acid boat conformation**. The files containing this data are provided.

| **System** | **k_on_** | | | **Probability** | | | **Total binding events** |
| --- | --- | --- | --- | --- | --- | --- | --- |
|  | **low** | **mean** | **high** | **low** | **mean** | **high** |  |
| **Primary site** | | | | | | | |
| Unglycosylated NA | 1.46/µM∙s | 1.63/µM∙s | 1.81/µM∙s | 3.24 x 10^-5^ | 3.62 x 10^-5^ | 4.00 x 10^-5^ | 362 |
| NA with Glyprot glycans | 1.76/µM∙s | 1.95/µM∙s | 2.14/µM∙s | 3.93 x 10^-5^ | 4.35 x 10^-5^ | 4.77 x 10^-5^ | 435 |
| NA with MD conf1 glycans | 0.12/µM∙s | 0.18/µM∙s | 0.24/µM∙s | 0.27 x 10^-5^ | 0.39 x 10^-5^ | 0.51 x 10^-5^ | 39 |
| NA with MD conf2 glycans | 0.03/µM∙s | 0.06/µM∙s | 0.10/µM∙s | 0.07 x 10^-5^ | 0.14 x 10^-5^ | 0.21 x 10^-5^ | 14 |
| NA with MD conf3 glycans | 1.11/µM∙s | 1.26/µM∙s | 1.41/µM∙s | 2.46 x 10^-5^ | 2.79 x 10^-5^ | 3.12 x 10^-5^ | 279 |
| **Secondary site** | | | | | | | |
| Unglycosylated NA | 0.24/µM∙s | 0.32/µM∙s | 0.40/µM∙s | 0.54 x 10^-5^ | 0.71 x 10^-5^ | 0.88 x 10^-5^ | 71 |
| NA with Glyprot glycans | 0.14/µM∙s | 0.20/µM∙s | 0.26/µM∙s | 0.31 x 10^-5^ | 0.45 x 10^-5^ | 0.58 x 10^-5^ | 45 |
| NA with MD conf1 glycans | 0.49/µM∙s | 0.60/µM∙s | 0.70/µM∙s | 1.06 x 10^-5^ | 1.29 x 10^-5^ | 1.52 x 10^-5^ | 129 |
| NA with MD conf2 glycans | 0.06/µM∙s | 0.10/µM∙s | 0.14/µM∙s | 0.13 x 10^-5^ | 0.22 x 10^-5^ | 0.31 x 10^-5^ | 22 |
| NA with MD conf3 glycans | 0.11/µM∙s | 0.17/µM∙s | 0.22/µM∙s | 0.25 x 10^-5^ | 0.37 x 10^-5^ | 0.49 x 10^-5^ | 37 |
| **Combined sites** | | | | | | | |
| Unglycosylated NA | 1.47/µM∙s | 1.64/µM∙s | 1.81/µM∙s | 3.26 x 10^-5^ | 3.64 x 10^-5^ | 4.02 x 10^-5^ | 364 |
| NA with Glyprot glycans | 1.88/µM∙s | 2.07/µM∙s | 2.26/µM∙s | 4.18 x 10^-5^ | 4.61 x 10^-5^ | 5.04 x 10^-5^ | 461 |
| NA with MD conf1 glycans | 0.67/µM∙s | 0.79/µM∙s | 0.91/µM∙s | 1.44 x 10^-5^ | 1.70 x 10^-5^ | 1.96 x 10^-5^ | 170 |
| NA with MD conf2 glycans | 0.08/µM∙s | 0.13/µM∙s | 0.18/µM∙s | 0.18 x 10^-5^ | 0.29 x 10^-5^ | 0.40 x 10^-5^ | 29 |
| NA with MD conf3 glycans | 1.25/µM∙s | 1.40/µM∙s | 1.56/µM∙s | 2.74 x 10^-5^ | 3.10 x 10^-5^ | 3.45 x 10^-5^ | 310 |

**Table S13**. **Association rate data for sialic acid chair conformation**. The files containing this data are provided.

| **System** | **k_on_** | | | **Probability** | | | **Total binding events** |
| --- | --- | --- | --- | --- | --- | --- | --- |
|  | **low** | **mean** | **high** | **low** | **mean** | **high** |  |
| **Primary site** | | | | | | | |
| Unglycosylated NA | 1.68/µM∙s | 1.87/µM∙s | 2.05/µM∙s | 3.69 x 10^-5^ | 4.09 x 10^-5^ | 4.49 x 10^-5^ | 409 |
| NA with Glyprot glycans | 1.84/µM∙s | 2.04/µM∙s | 2.23/µM∙s | 4.07 x 10^-5^ | 4.49 x 10^-5^ | 4.91 x 10^-5^ | 449 |
| NA with MD conf1 glycans | 0.22/µM∙s | 0.29/µM∙s | 0.36/µM∙s | 0.46 x 10^-5^ | 0.62 x 10^-5^ | 0.78 x 10^-5^ | 62 |
| NA with MD conf2 glycans | 0.02/µM∙s | 0.06/µM∙s | 0.09/µM∙s | 0.05 x 10^-5^ | 0.12 x 10^-5^ | 0.19 x 10^-5^ | 12 |
| NA with MD conf3 glycans | 1.19/µM∙s | 1.34/µM∙s | 1.50/µM∙s | 2.60 x 10^-5^ | 2.94 x 10^-5^ | 3.28 x 10^-5^ | 294 |
| **Secondary site** | | | | | | | |
| Unglycosylated NA | 0.00/µM∙s | 0.00/µM∙s | 0.00/µM∙s | 0.00 x 10^-5^ | 0.00 x 10^-5^ | 0.00 x 10^-5^ | 0 |
| NA with Glyprot glycans | 0.00/µM∙s | 0.00/µM∙s | 0.00/µM∙s | 0.00 x 10^-5^ | 0.00 x 10^-5^ | 0.00 x 10^-5^ | 0 |
| NA with MD conf1 glycans | 0.00/µM∙s | 0.00/µM∙s | 0.00/µM∙s | 0.00 x 10^-5^ | 0.00 x 10^-5^ | 0.00 x 10^-5^ | 0 |
| NA with MD conf2 glycans | 0.00/µM∙s | 0.00/µM∙s | 0.00/µM∙s | 0.00 x 10^-5^ | 0.00 x 10^-5^ | 0.00 x 10^-5^ | 0 |
| NA with MD conf3 glycans | 0.00/µM∙s | 0.00/µM∙s | 0.00/µM∙s | 0.00 x 10^-5^ | 0.00 x 10^-5^ | 0.00 x 10^-5^ | 0 |
| **Combined sites** | | | | | | | |
| Unglycosylated NA | 1.75/µM∙s | 1.93/µM∙s | 2.13/µM∙s | 3.84 x 10^-5^ | 4.25 x 10^-5^ | 4.66 x 10^-5^ | 425 |
| NA with Glyprot glycans | 1.87/µM∙s | 2.06/µM∙s | 2.26/µM∙s | 4.11 x 10^-5^ | 4.54 x 10^-5^ | 4.97 x 10^-5^ | 454 |
| NA with MD conf1 glycans | 0.20/µM∙s | 0.27/µM∙s | 0.34/µM∙s | 0.43 x 10^-5^ | 0.58 x 10^-5^ | 0.73 x 10^-5^ | 58 |
| NA with MD conf2 glycans | 0.01/µM∙s | 0.03/µM∙s | 0.06/µM∙s | 0.02 x 10^-5^ | 0.07 x 10^-5^ | 0.12 x 10^-5^ | 7 |
| NA with MD conf3 glycans | 1.27/µM∙s | 1.43/µM∙s | 1.59/µM∙s | 2.77 x 10^-5^ | 3.12 x 10^-5^ | 3.47 x 10^-5^ | 312 |
